# DNA replication timing directly regulates the frequency of oncogenic chromosomal translocations

**DOI:** 10.1101/2021.05.29.446276

**Authors:** Mihaela Peycheva, Tobias Neumann, Daniel Malzl, Mariia Nazarova, Ursula Schoeberl, Rushad Pavri

**Affiliations:** Research Institute of Molecular Pathology (IMP), Vienna BioCenter (VBC), Campus-Vienna-Biocenter 1, 1030 Vienna, Austria

**Keywords:** Translocations, DNA replication timing, antibody maturation, genome architecture

## Abstract

Chromosomal translocations result from the joining of DNA double-strand breaks (DSBs) and frequently cause cancer. Yet, the steps linking DSB formation to DSB ligation remain undeciphered. We report that DNA replication timing (RT), mediated by replication origin activity, directly regulates the genesis of lymphomagenic *Myc* translocations during antibody maturation in B cells. Reduced levels of the replicative helicase, the minichromosome-maintenance (MCM) complex, decreases translocations and globally abrogates the RT program. Ablating a single replication origin at *Myc* causes an early-to-late RT switch with major loss of translocations, a phenotype that is reversed by restoring early RT. Finally, this novel RT-regulated mechanism occurs after DSB formation and independently of DSB frequency. Thus, RT constitutes a distinct regulatory event in translocation biogenesis linking DSB formation to DSB ligation.

## Introduction

In antigen-activated B lymphocytes, deleterious translocations arise during the physiological process of antibody maturation when B cells diversify their antibody repertoire via somatic hypermutation and isotype class switch recombination at the immunoglobulin (Ig) genes (*1–6*). Importantly, antibody maturation involves DSB formation at Ig genes induced by the mutagenic action of activation induced deaminase (AID) in a transcription-dependent manner (*7–10*). Consequently, antibody maturation is inextricably linked to genome instability because AID also targets major transcribed oncogenes in B cells like *Myc* and *Bcl6*, resulting in DSBs and translocations between these oncogenes and the Ig loci, an event that deregulates oncogene expression and triggers tumorigenesis (*11–18*). *Myc* and Ig translocations are frequently observed in major mature B cell cancers, including diffuse large B cell lymphoma, multiple myeloma and, most strikingly, in Burkitt lymphoma where *Myc-Ig* translocations are seen in all tumors and are proposed to be the primary drivers of tumorigenesis (*19–21*). These translocations have been recapitulated not only in mouse tumor models (*22–24*) but, most pertinently, also in primary B cells (*25–29*) and the murine B lymphoma line, CH12 (*30, 31*), upon AID overexpression, making the latter systems ideal for addressing the mechanisms of translocation biogenesis. In this study, we retrovirally express AID fused to the estrogen receptor ligand-binding domain (AIDER) (*32*) in CH12 cells (*33*) or use primary splenic B cells expressing AIDER from the *Rosa26* locus (*23*), and add 4-hydroxy tamoxifen (4-HT) to trigger nuclear import of AIDER.

## Results

### The replicative helicase, the MCM complex, is required for oncogenic *Myc-Igh* translocations

It has been shown that reduced levels of the hexameric replicative helicase, the minichromosome maintenance (MCM) complex, can lead to cancer in mice resulting from increased replicative stress-mediated genome instability (*34, 35*). Therefore, we reasoned that MCM complexes may play a protective role in B cells during AID-mediated translocation genesis which, importantly, occurs in B cells undergoing massive antigen-mediated proliferation with very short cell cycles (6-8 h) (*36*). We depleted Mcm6 with a short hairpin RNA (shRNA) (henceforth called shMcm6) in primary, activated splenic B cells and in activated CH12 cells. Under these conditions, shMcm6 cells express ~20% of Mcm6 protein compared to control cells (shLacZ) (Fig. S1A), are proliferative (Fig. S1B) and show normal DNA synthesis (Fig. S1C). We note that the normal growth and viability of MCM-depleted cells is consistent with several previous reports in a variety of cell types (*37–41*). Total RNA sequencing (RNA-seq) showed that Mcm6 RNA, but not other MCM subunits, was strongly downregulated in shMcm6 cells (Fig. 1A and Fig. S1 D, E). In contrast, nuclear mass spectrometry revealed decreased protein expression of Mcm6 (~3.5-fold) as well as other MCM complex subunits (Fig. 1B and Fig. S1F). Thus, the MCM complex is destabilized at the protein level when Mcm6 protein is limiting. Importantly, we did not detect significant loss of other DNA replication factors or DNA repair factors, both at the RNA and protein levels (Fig. 1A, B). Also, shMcm6 cells do not show major changes in RNA or protein expression of AID, *Myc* and Ig genes (Fig. S1 E, F), or expression of AID translocation hotspot genes (Fig. 1A).

**Figure 1.**
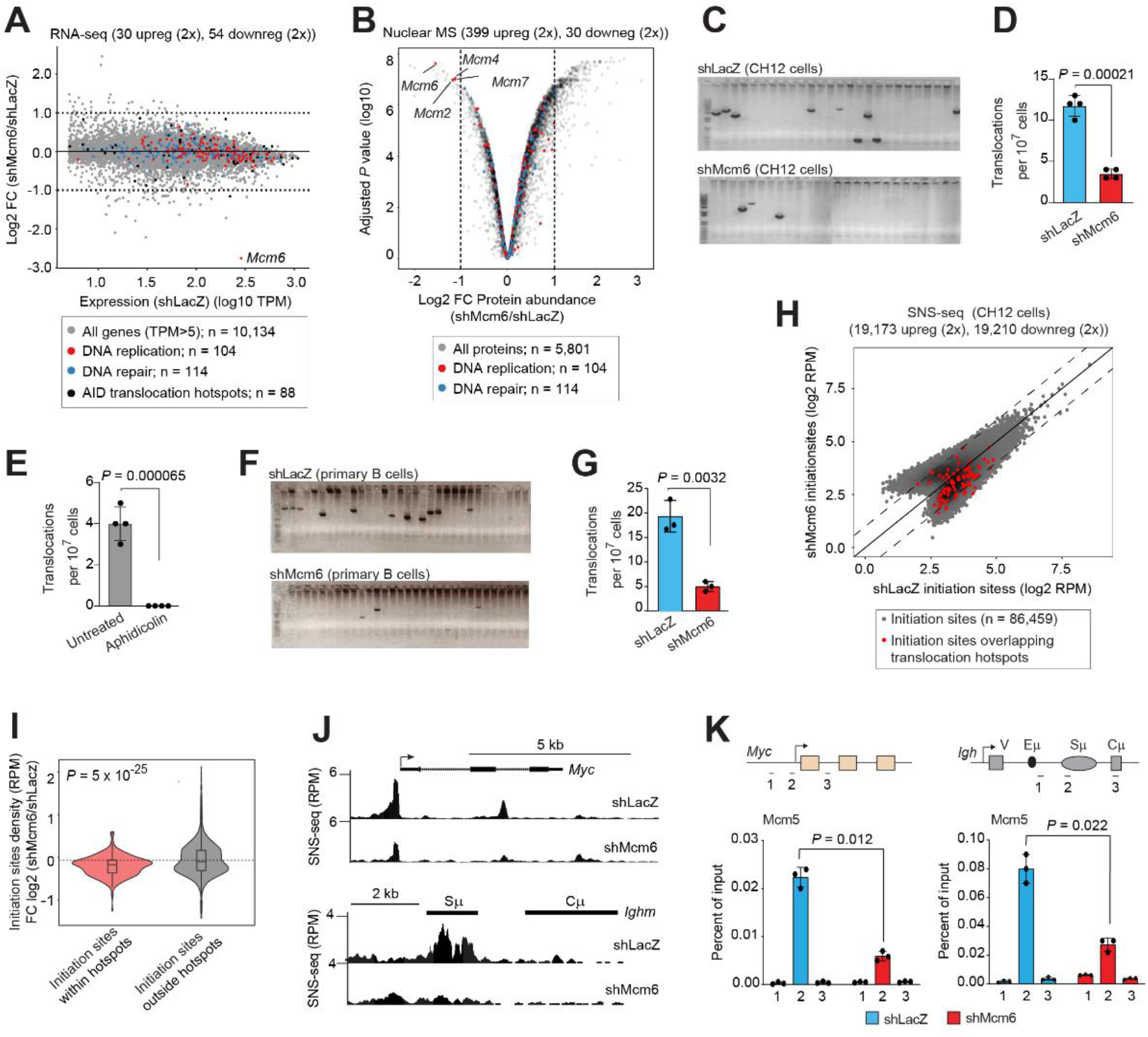
AID-dependent *Myc*-*Igh* translocation frequency correlates with the frequency of replication origin activity at *Myc* and *Igh*. **(A)** RNA-seq analysis in shLacZ or shMcm6 CH12 cells (averaged from triplicate experiments) and represented as an MA plot of fold change (shMcm6/shLacZ) against RNA expression in shLacZ cells calculated as transcripts per million (TPM). Dots represent mRNAs with TPM>5, a cutoff for expressed genes. The dotted lines demarcate 2-fold upregulated (log2 FC > 1) or 2-fold downregulated (log2 FC < −1) RNAs. Red, blue and black dots represent DNA replication genes, DNA repair genes and AID translocation hotspots, respectively. The location of Mcm6 RNA is indicated. **(B)** Nuclear mass spectrometric analysis from shLacZ and shMcm6 CH12 cells represented as a Volcano plot of fold-change (shMcm6/shLacZ) against the adjusted *P* value. The dotted lines demarcate 2-fold upregulated (log2 FC > 1) or 2-fold downregulated (log2 FC < −1) proteins. **(C)** A representative agarose gel revealing translocation PCR products from genomic DNA of activated CH12 cells expressing AIDER and infected with lentiviruses expressing shRNAs against LacZ (shLacZ) or Mcm6 (shMcm6). 4-HT was added at the time of activation to induce nuclear import of AIDER. **(D)** Quantitation of translocations following Sanger sequencing of PCR products. The data represent four independent experiments. **(E)** *Myc-Igh* translocation assay as above in WT activated CH12 cells expressing AIDER treated with Aphidicolin (1mM) for 48 hr. **(F)** Translocation PCR assay from primary, activated splenic B cells expressing AIDER from the *Rosa26* locus, infected with retroviruses expressing shLacZ or shMcm6, and treated with 4-HT. **(G)** Quantification from three independent translocation PCR experiments in primary, splenic B cells. **(H)** SNS-seq analysis in shLacZ and shMcm6 CH12 cells. Each dot represents a replication initiation site identified via peak calling and overlapping with initiation zones (see also Fig. S2C). Red dots are replication initiation sites overlapping AID-dependent translocation hotspots (Supplementary Table 1). Initiation site density is calculated as reads per million (RPM). The dotted lines indicate 2-fold change on either side of the diagonal. **(I)** Violin plots based on data from **(H)** showing the highly significant enrichment of replication initiation sites within AID translocation hotspots compared to initiation sites outside AID translocation hotspots. **(J)** UCSC browser snapshots of SNS-seq profiles at *Myc* (top) and *Igh* (bottom) from shLacZ and shMcm6 cell. **(K)** ChIP-qPCR for Mcm5 occupancy at *Myc* (left) and *Igh* (right) in shLacZ and shMcm6 CH12 cells. The location of amplicons (1–3) is indicated in the upper panel. Data represent three independent experiments. All *P* values in the above figures were determined by the unpaired Student’s t test.

Translocations between *Myc* and the Ig heavy chain (*Igh*) locus in AIDER-expressing shLacZ and shMcm6 cells were quantified using a previously established nested PCR assay (*29*) (Fig. S1G). Contrary to our expectation, we observed a significant decrease in *Myc-Igh* translocation frequency in shMcm6 CH12 cells relative to the shLacZ control (Fig. 1C, D). Moreover, these translocations were almost completely abrogated when wild-type (WT) cells were arrested at the G1/S phase boundary with the DNA polymerase inhibitor, Aphidicolin (Fig. 1E). *Myc-Igh* translocations were also strongly reduced in shMcm6 primary splenic B cells (Fig. 1F, G). We conclude that, in contrast to studies in other systems (*34, 35*), MCM depletion in activated B cells suppresses genome instability during antibody maturation. Moreover, this function of the MCM complex is exerted downstream of AID targeting as seen by the normal rates of AID-mediated mutation at *Myc* and *Igh* in shMcm6 cells (Fig. S1 H, I). In sum, these results reveal a novel role of the MCM complex as a driver of B cell lymphomagenesis.

### *Myc*-*Igh* translocation frequency correlates with the activity of origins of replication at *Myc* and *Igh*

The loading of MCM complexes to origins of replication licenses these origins for activation by replication firing factors during S phase (*42*). Importantly, the normal cycling of shMcm6 cells (*43*) implied that the act of DNA replication was unlikely to be a causal factor in driving *Myc-Igh* translocations. Rather, we hypothesized that MCM depletion would reduce the licensing of origins and thereby decrease origin activation frequencies at AID target genes. Therefore, we performed short nascent strand sequencing (SNS-seq) in CH12 cells which measures the relative enrichment of nascent leading strands at origins of replication (*44, 45*) (Fig. S2A). In effect, replication initiation sites identified by SNS-seq enrichments reflect the relative firing frequency of the underlying replication origins in the population. Two independent SNS-seq experiments were performed in CH12 cells, each with two replicates (Fig. S2B). Only replication initiation sites that reproducibly fell within initiation zones were retained (Fig. S2C).

We observed a significant alteration of the replication origin landscape in shMcm6 cells marked by several upregulated and downregulated replication initiation sites relative to shLacZ cells (Fig. 1H). By overlapping initiation sites with selected chromatin marks and genomic compartments (obtained from Hi-C), we observed that downregulated initiation sites were located mostly in active chromatin within A compartments whereas upregulated initiation sites overlapped largely with histone H3 trimethylated at lysine 9 (H3K9me3)-rich constitutive heterochromatin within B compartments (Fig. S2 D, E). Using a statistical permutation test, we found that AID-dependent translocation hotspots overlap with replication initiation sites significantly more often than compared to a null distribution of randomly placed translocation hotspots and initiation sites (Fig. S2F). Most importantly, initiation sites overlapping AID-dependent translocation hotspots were mostly downregulated in shMcm6 cells and with high significance (Fig. 1H, I). Thus, AID-dependent translocation hotspots are enriched in MCM-sensitive replication initiation zones.

At *Myc*, we observed a strong origin of replication just upstream of the promoter that was downregulated in shMcm6 cells (Fig. 1J, upper panel). SNS-qPCR analysis confirmed this decrease in origin activity in CH12 cells as well as in primary splenic B cells (Fig. S2G). At *Igh*, SNSs were enriched within the switch recombination sequences and these origins were downregulated in shMcm6 cells (Fig. 1J, lower panel). We confirmed the loss of origin activity at *Igh* with SNS-qPCR in both CH12 and primary B cells (Fig. S2H). Finally, at both *Myc* and *Igh*, the decrease in origin activity correlated with significantly reduced MCM complex occupancy indicative of decreased origin licensing (Fig. 1K). We conclude that the frequency of *Myc-Igh* translocation correlates with the frequency of replication origin licensing and origin activity at *Myc* and *Igh*.

### Replication origin activity at *Myc* directly regulates *Myc*-*Igh* translocation frequency

To determine whether the observed correlation between translocation frequency and replication origin activity reflects a direct, causal relationship between these processes, we deleted the promoter-proximal origin of replication at *Myc* in CH12 cells, spanning 800 bp, with CRISPR editing (Fig. 2A upper panel, and Fig. S3A). We hypothesized that if replication origin activity was causal for translocation genesis, then deletion of the origin of replication would impair translocation frequency independently of *Myc* transcription and AID-mediated mutagenesis. Importantly, the deletion does not overlap the core promoter, as determined by the occupancy of TBP and TFIIB, core components of the transcription preinitiation complex (Fig. 2A). In addition, the deletion does not overlap with the major transcription initiation sites identified by precision run-on sequencing for capped transcripts (PRO-cap) (*46*), which maps the location and orientation of capped transcription initiation sites associated with actively engaged RNA polymerase II (Fig. 2A).

**Fig 2.**
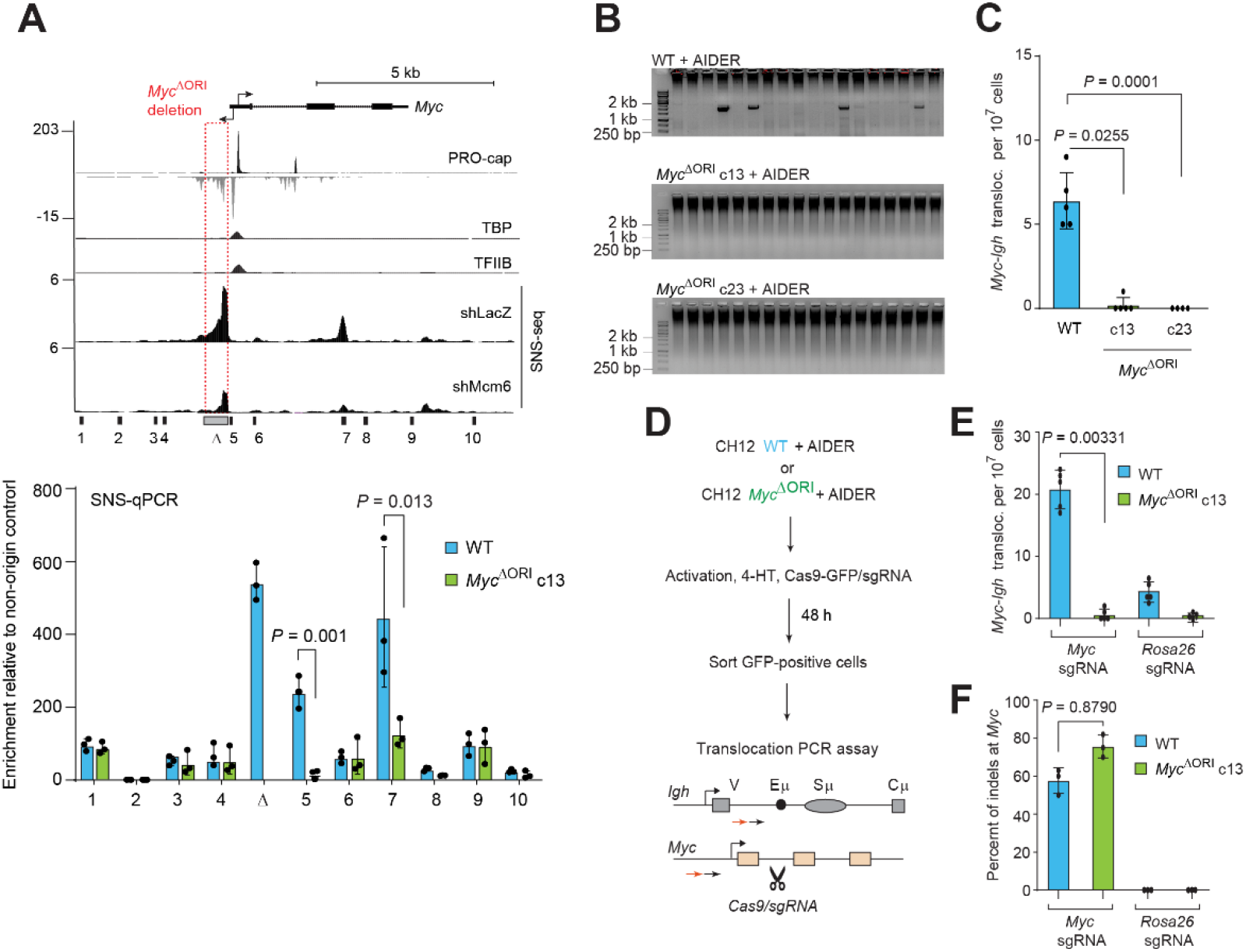
Deletion of a single origin of replication at *Myc* strongly decreases *Myc-Igh* translocation rate downstream of DSBs and independent of DSB frequency. **(A)** <u>Top pane</u>l: Snapshot of the *Myc* locus showing the SNS-seq profiles in shLacZ and shMcm6 cells. Also shown are CH12 (WT) tracks of PRO-cap (indicating capped transcription initiation sites), TBP and TFIIB (both indicating the core promoter). The origin targeted for CRISPR-mediated deletion is boxed in red and lies upstream of both the core promoter and the major antisense initiation site. <u>Bottom pane</u>l: SNS-qPCR at the *Myc* locus in WT and *Myc*^ΔORI^ clone 13 (c13) cells. Amplicons for qPCR are shown in the upper panel below the SNS-seq snapshot. Enrichments are calculated relative to a non-origin control. **(B)** Representative agarose gel of a *Myc-Igh* translocation PCR assay in WT, *Myc*^ΔORI^ c13 and *Myc*^ΔORI^ c23 cells following AIDER expression and 4-HT addition. **(C)** Quantification of results from five independent *Myc-Igh* translocation PCR assays in WT, *Myc*^ΔORI^ c13 and *Myc*^ΔORI^ c23 cells. **(D)** Scheme of *Myc-Igh* translocation PCR assay in WT and *Myc*^*ΔORI*^ c13 cells expressing AIDER and transfected with Cas9 and a *Myc*-specific sgRNA to bypass DSB formation at *Myc*. **(E)** Quantification of *Myc-Igh* translocation PCR assay following Cas9/sgRNA DSB induction at *Myc* in AIDER-expressing WT and *Myc*^ΔORI^ c13 cells from four independent replicates. The *Rosa26*-specific sgRNA was used as a control to compare with the effect of the *Myc* sgRNA. **(F)** Sanger sequencing analysis of the region flanking the sgRNA target sequence at *Myc* represented as the percentage of insertions or deletions (indels) observed at the Cas9 cleavage site. All *P* values in the figure were determined by the unpaired Student’s t test.

Two independent deletion clones (c13 and c23) were chosen for analysis and henceforth called *Myc*^ΔORI^ c13 and *Myc*^ΔORI^ c23 (Fig. S3 B, C). Both clones were viable and neither suffered from proliferation defects (Fig. S3D). Total RNA-seq revealed no major changes in expression of AID target genes or DNA replication and repair genes (Fig. S3E). Importantly, *Myc* itself was expressed at similar levels in both *Myc*^ΔORI^ clones (Fig. S3F). Crucially, class switch recombination was robust in both *Myc*^ΔORI^ lines, indicating that transcription-coupled mutagenesis, DSB formation, DNA repair and non-homologous end joining pathways, all of which are essential for class switch recombination and translocations, were fully functional in these clones (Fig. S3 G, H). Furthermore, AID-mediated mutation frequencies at *Myc* in *Myc*^ΔORI^ cells were comparable to WT cells implying that AID targeting is unaffected by the loss of replication origin activity (Fig. S3I). Therefore, the minor gene expression differences observed between the two *Myc*^ΔORI^ lines are likely due to their different clonal origins and have no significant impact on major physiological processes in these cells.

SNS-qPCR analysis confirmed the loss of origin activity at the deleted region in *Myc*^ΔORI^ cells (Fig. 2A, bottom panel bar graph). However, *Myc*^ΔORI^ cells also showed a decrease in the activity of the intragenic origin of replication, mimicking the results from shMcm6 cells where this intragenic origin was also downregulated (compare SNS-seq tracks with the SNS-qPCR bar graphs in Fig. 2A). Consequently, the replication origin landscape at the *Myc*^ΔORI^ allele resembles that of *Myc* in shMcm6 cells. Most importantly, *Myc-Igh* translocation frequency was severely reduced in *Myc*^ΔORI^ cells relative to WT cells (Fig. 2 B, C). Thus, replication origin activity at *Myc* has a direct and profound effect on the genesis of *Myc-Igh* translocations downstream of AID-mediated mutagenesis..

### DNA replication origin activity regulates a distinct step in translocation genesis downstream of DSB formation and independent of DSB frequency

Given that AID targeting was unaffected in *Myc*^ΔORI^ cells (Fig. S3I), we asked whether DSB formation, the next obligatory step for translocations, was impacted. Due to the low mutation rate coupled with the low density of AID hotspot motifs at non-*Ig* AID targets like *Myc*, DSBs occur rarely and are challenging to detect. However, we reasoned that if DSB formation was indeed limiting in *Myc*^ΔORI^ cells, then a potent exogenous DSB within the translocation hotspot should rescue the *Myc-Igh* translocation rate. Therefore, we transfected AIDER-expressing *Myc*^ΔORI^ cells with Cas9 and a small guide RNA (sgRNA) targeting the first intron of *Myc*, a region that lies within the known translocation hotspot (*30*). Cas9/sgRNA-transfected WT and *Myc*^ΔORI^ cells were sorted and analyzed for *Myc-Igh* translocations (Fig. 2D). Surprisingly, we detected virtually no increase in *Myc-Igh* translocation frequency in *Myc* sgRNA-expressing *Myc*^ΔORI^ cells relative to cells with a control sgRNA (Fig. 2E). This was not due to inefficient Cas9 activity since 70-80% of WT and *Myc*^ΔORI^ cells harbored indels at the Cas9 cleavage site (Fig. 2F). We therefore conclude that the *Myc* origin regulates translocation frequency downstream of DSBs. Moreover, since the frequency of the Cas9-induced DSB is orders of magnitude greater than that of a physiological AID-induced DSB, we conclude that the *Myc* origin regulates translocation genesis completely independently of DSB frequency.

To determine whether other AID-dependent translocations are also impacted in *Myc*^ΔORI^ cells, we performed a modified version of linear amplification-mediated high-throughput genome-wide translocation sequencing (LAM-HTGTS (*47*); Methods) which employs an exogenously introduced bait DSB to identify associated prey translocations by locus-specific enrichment and deep sequencing. A high-frequency bait DSB was introduced in *Myc* with Cas9 and a *Myc*-specific sgRNA (*30*). This procedure inherently bypasses the AID-dependent DSB formation step at *Myc*, thereby allowing us to determine, on a genome-wide scale, whether decreased replication origin activity at *Myc* (*Myc*^ΔORI^ cells) or AID target genes generally (shMcm6 cells) regulates *Myc* translocation genesis upstream or downstream of DSB formation.

We performed LAM-HTGTS in AIDER-expressing WT, *Myc*^ΔORI^ c13 (Fig. 3A) and *Myc*^ΔORI^ c23 (Fig. 3B), as well as in shLacZ and shMcm6 cells (Fig. 3C). In this system, the DSB at *Myc* is generated primarily by Cas9 whereas AIDER is required to ensure high rates of physiological DSBs at other AID target genes. Three independent LAM-HTGTS experiments were conducted and libraries were sequenced with the MinIon flow cell (Oxford Nanopore Technologies), which detects translocation breakpoints at single-nucleotide resolution by spanning the entire site with long reads. Raw translocations involving the *<u>Myc</u>* DSB were seen in all conditions indicating that translocation genesis was not impaired per se in these cells (Fig. S3 J to L) From these data, we defined 88 high-confidence AID-dependent hotspots (Supplementary Table 1) based on the density of translocations, consistency between replicates and several quality criteria (Methods). Strikingly, decreased translocation frequency was observed at virtually all AID-dependent hotspots in shMcm6 cells and both *Myc*^ΔORI^ lines as seen from the circos plots showing fold-changes in translocations (Fig. 3 A to C) and scatter plots of translocation hotspot densities (Fig. 3 D to F).

**Fig 3:**
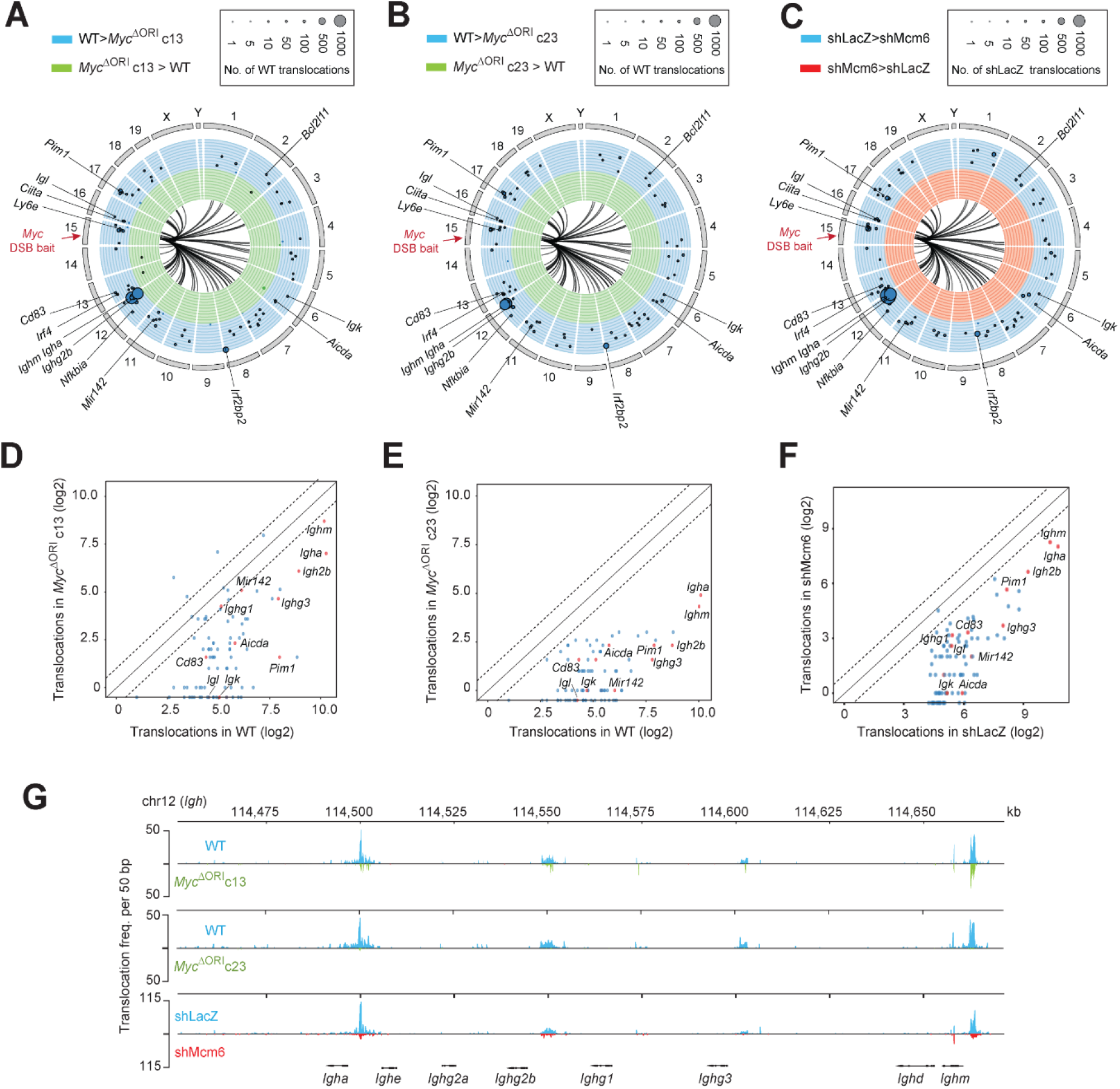
AID-dependent translocations are globally reduced in *Myc*^ΔORI^ and shMcm6 cells. **(A) to (C)** Circos plots displaying AID-dependent translocations obtained from linear amplification-mediated high-throughput genome-wide translocation sequencing (LAM-HTGTS) in *Myc*^ΔORI^ c13 vs WT cells (A), *Myc*^ΔORI^ c23 vs WT cells (B) and shMcm6 vs shLacZ cells (C). The location of the Cas9-induced *Myc* bait DSB is indicated in red text. Each arc radiating from *Myc* indicates an AID-dependent *Myc* translocation hotspot to the indicated chromosome. The plots show the fold-change of translocations (log2 WT/ *Myc*^ΔORI^ in (A) and (B) or log2 shMcm6/shLacZ in (C) in the following manner: each plot is divided into concentric circles ranging from lowest fold-change (log2 FC = −7, innermost circle) to highest fold-change (log2 FC = 7, outermost circle) with the middle circle (the border between the two colors) indicating no change (log2 FC = 0). Each dot on the plot represents a hotspot and the dot size correlates with the raw numbers of translocations within that hotspot in WT cells, as shown in the key above each plot. Thus, for example, the *Igh* locus has the largest dot size in (A) to (C) indicating that it is the most frequent translocation partner of *Myc* in WT and shLacZ cells, as expected. Dots in the blue portion of the plot represent hotspots where translocation frequency is reduced in *Myc*^ΔORI^ or shMcm6 cells (C) whereas dots in the green portion represent hotspots with increased translocation frequency in *Myc*^ΔORI^ or shMcm6 cells. The presence of mostly blue dots indicates that translocation frequency is reduced at nearly all hotspots in *Myc*^ΔORI^ and shMcm6 cells. Selected AID translocation hotspots found recurrently in many studies are highlighted. **(D) to (F)** Scatter dot plots showing the quantification of translocation frequency in *Myc*^ΔORI^ c13 vs WT cells (D), *Myc*^ΔORI^ c23 vs WT cells (E) and shMcm6 vs shLacZ cells (F). Selected hotspots (same as in (A) to (C)) are indicated as red dots. The dotted lines indicate two-fold change (log2) on either side of the diagonal. **(G)** Snapshot of *Myc* translocations to the *Igh* locus in the indicated conditions. The location of the various *Igh* constant region genes is indicated at the bottom. *Ighm, Igh2b* and *Igha* are transcribed in these cells and hence harbor the vast majority of translocations

The transcribed *Igh* genes (*Ighm, Igha* and *Igh2b*) were the strongest hotspots in WT and shLacZ cells, as expected given that they are the prime targets of AID in activated CH12 cells (Fig. 3 A to F). Moreover, translocations to all these genes were strongly reduced in all conditions (Fig. 3 A to G). In addition, WT and shLacZ cells harbored several known and recurrent AID-dependent hotspots, including *Igk* and *Igl* (encoding the Ig*κ* and Ig*λ* antibody light chains), *Cd83, Pim1, Mir142 and Aicda* (Fig. 3 A to F and Fig. S4, S5, S6) (*26, 27, 30, 48*). Examples of LAM-HTGTS results at various non-*Igh* hotspots are provided in Fig. S4 (WT versus *Myc*^ΔORI^ c13), Fig. S5 (WT versus *Myc*^ΔORI^ c23) and Fig. S6 (shLacZ versus Mcm6).

We conclude that replication origin activity at *Myc* directly regulates genome-wide AID-dependent translocation frequency at a step downstream of DSB formation and independently of DSB frequency. Moreover, we infer that the translocation defect in shMcm6 cells arises due to decreased local origin activity at *Myc* and other AID target genes (Fig. 1 F to I) because the translocation phenotype of *Myc*^ΔORI^ cells, where replication origin activity is locally decreased at *Myc*, is virtually identical to that of shMcm6 cells, where replication origin activity of AID targets is globally decreased.

### DNA replication timing (RT) is locally perturbed at *Myc* in *Myc*^ΔORI^ cells and globally abrogated in shMcm6 cells

Having established that local replication origin activity at *Myc* has a direct and significant role in the genesis of *Myc* translocations, we investigated the underlying mechanism. Replication origin activation has been proposed to occur in clusters or hubs, in which neighboring origins in 3D space are colocalized and fire synchronously (*49, 50*). Because this event involves the spatial proximity of replication origins, we hypothesized that shared replication timing (RT) of neighboring origins in AID target genes would also lead to the colocalization of DSBs in the vicinity of these origins. This hypothesis predicts that: (1) AID target genes have similar RT in WT cells, (2) shMcm6 cells have altered RT of at least the AID target genes, and (3) that the *Myc*^ΔORI^allele has an altered RT compared to the WT *Myc* allele.

To measure RT, we performed Repli-seq in shMcm6 and *Myc*^ΔORI^ cells. Repli-seq involves pulse-labeling of S phase cells via incorporation of 5-bromo-2’-deoxyuridine (BrdU) into replicating DNA followed by isolation of early (E) and late (L) S phase cells, genomic DNA extraction, enrichment of BrdU-labeled DNA and quantification of enriched DNA with deep sequencing (Fig. S7 A, B) (*51, 52*). WT and shLacZ cells showed distinct and largely non-overlapping early and late domains (Fig. 4 A, B). RT is displayed as a log2 E/L ratio where early and late domains are demarcated by positive and negative values, respectively (Fig. 4 A, B). This is also visualized in metaplots showing a wide separation between early and late replication domains in WT and shLacZ cells (Fig. 4C). *Myc*^ΔORI^ cells showed similar profiles to WT, although the metaplots revealed a slight shift in the early RT peak relative to WT cells (Fig. 4C), which we infer to be due to clonal variation between the *Myc*^ΔORI^ and the parental WT cells. Thus, we conclude that *Myc*^ΔORI^ cells do not undergo global changes in RT.

**Fig. 4.**
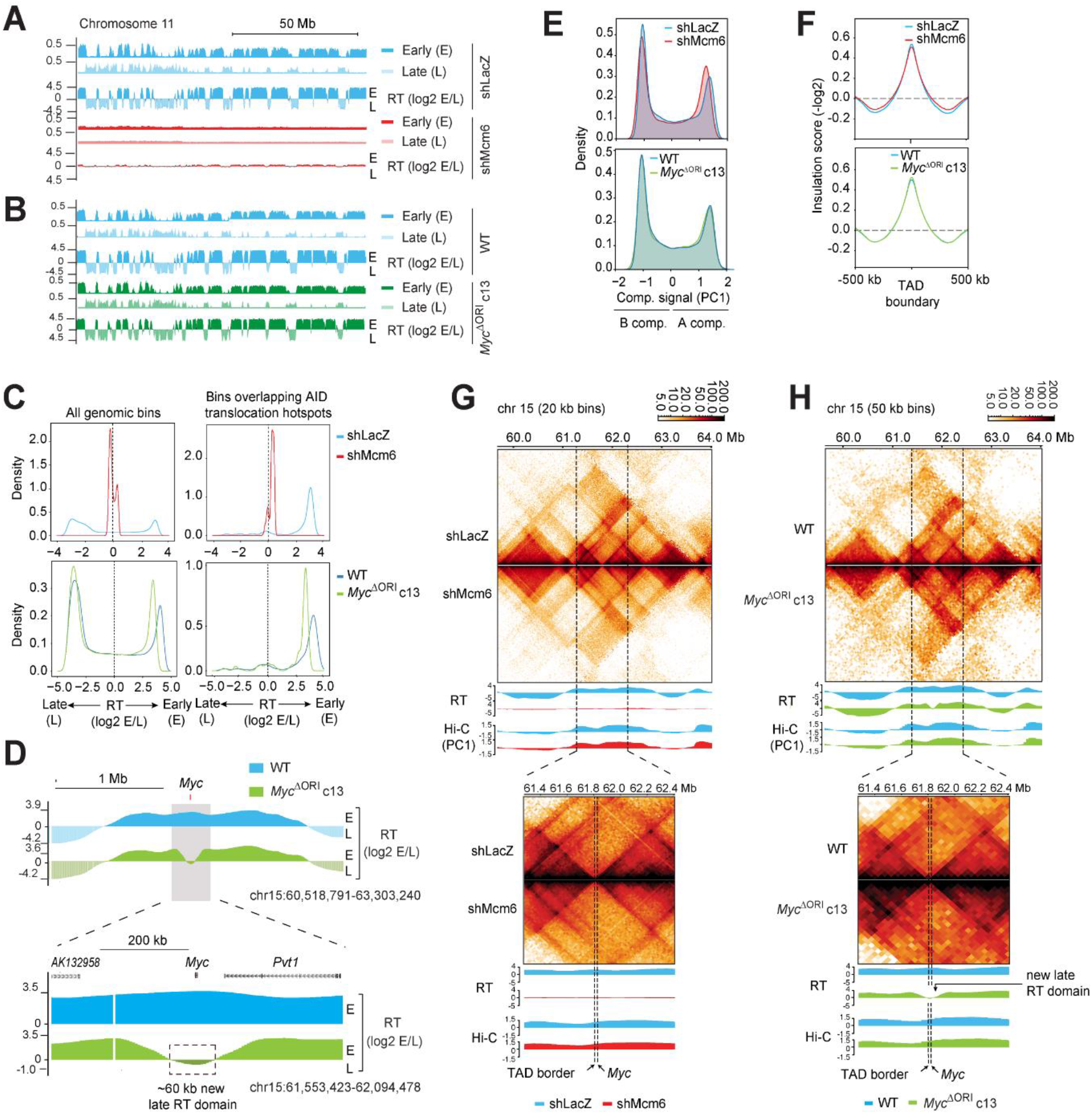
Replication timing (RT) is globally disrupted in shMcm6 cells and locally at *Myc* in *Myc*^ΔORI^ cells. **(A)** Snapshot of the whole chromosome 11 showing the individual early (E) and late (L) Repli-seq tracks followed by the RT (log2 E/L ratio) tracks for shLacZ and shMcm6 where positive and negative values reflect early and late RT, respectively. **(B)** Snapshot of chromosome 11 showing E, L, and RT tracks from WT and *Myc*^ΔORI^ c13 cells. **(C)** Repli-seq derived metaplots showing the RT profiles (log2 E/L ratio) upon Mcm6 depletion (top) and in *Myc*^ΔORI^ c13 cells (bottom). The left plots show composite genome-wide profiles and right plots display profiles of only AID-dependent translocation hotspots identified from LAM-HTGTS in Fig. 3. **(D)** RT tracks at the *Myc* locus in WT and *Myc*^ΔORI^ cells showing the large 1.8 Mb early-replicating domain in WT cells (top) and a zoomed-in view (bottom) highlighting the new ~60 kb late RT domain created in *Myc*^ΔORI^ cells (boxed). **(E)** Histogram of Hi-C genome compartmentalization quantified as the first principal component (PC1) from the Hi-C matrix where positive and negative values reflect A and B compartments, respectively. **(F)** Average insulation score around the boundaries of 553 TADs (400-900 kb) computed from a 10 kb KR-normalized contact matrix. **(G)** Hi-C matrices at *Myc* in shLacZ and shMcm6 cells showing an ~5 Mb region with several TADs (top panel) and a zoomed-in ~1 Mb region (bottom panel) showing two TADs with *Myc* lying in the right TAD ~6 kb from the TAD border. Below each matrix are the Repli-seq RT tracks and genome compartmentalization tracks (Hi-C PC1). Note how the PC1 profile is unchanged in shMcm6 cells whereas RT is strongly attenuated. **(H)** Hi-C matrices at *Myc* displayed exactly as in (G) from WT and *Myc*^ΔORI^ c13 cells. Note how the shift to late RT at *Myc* is not accompanied by a change in compartmentalization. Note also that TAD integrity is intact and that the new RT domain created in *Myc*^ΔORI^ cells spans well across the TAD border.

In shMcm6 cells, however, a dramatic loss of all early and late domains was observed (note the raw E and L tracks in Fig. 4A). This resulted in log2 E/L values close to zero manifesting as a severely flattened RT profile across the length of all chromosomes (Fig. 4 A to C). Thus, shMcm6 cells appear to undergo temporally uncoordinated replication in which any origin is capable of firing at any time in S phase. This complete abrogation of the RT program reveals that the endogenous levels of MCM complexes are absolutely critical for establishing a normal RT program. These findings demonstrate that changes in origin activity directly and profoundly impact on RT.

Importantly, AID-dependent translocation hotspots (Fig. 3) have a highly positive log2 E/L ratio in WT and shLacZ cells, indicating that they are mostly early-replicating (Fig. 4C). This profile was unchanged in *Myc*^ΔORI^ cells, indicating that these cells retain early replication of AID targets (Fig. 4C). In contrast, in shMcm6 cells, these hotspots have log2 E/L ratios close to zero, similar to the average genomic profile, indicating that AID targets, like the rest of the genome, have lost their early RT potential in shMcm6 cells and now replicate in an uncoordinated manner throughout S phase (Fig. 4C).

In contrast to the unchanged RT of other AID targets in *Myc*^ΔORI^ cells, the RT profile at *Myc* in *Myc*^ΔORI^ cells was markedly altered. *Myc* is located within a ~1.8 Mb early-replicating domain (Fig. 4D, top panel). Strikingly, at the *Myc*^ΔORI^ allele, a new ~250 kb domain was created that is characterized by a continuous decrease in early RT relative to the WT allele with the middle ~60 kb portion, which harbors *Myc*, exhibiting late replication (Fig. 4D; bottom panel zoomed-in view). The switch in RT at *Myc* was confirmed by Repli-qPCR analysis (Fig. S7C). These data suggest that this new replication domain is passively replicated by replication forks originating from neighboring early RT domains in *Myc*^ΔORI^ cells. We therefore conclude that the activity of the origin of replication at *Myc* is essential to establish early RT. Along with the results from shMcm6 cells, these data demonstrate that the activity of origins of replication can have a profound impact on RT. Thus, early RT of AID hotpots is tightly linked to the loss of translocations in both shMcm6 cells and *Myc*^ΔORI^ cells.

### Changes in RT are not associated with altered genome architecture

RT is strongly correlated with genome compartmentalization, namely, active A and silent B compartments typically replicate early and late, respectively (*53*). The fact that compartmentalization and RT are established concurrently following exit from mitosis has led to the proposal that these two features are mechanistically linked (*50*). Given these observations, we performed Hi-C (*54*) to determine whether the changes in RT were accompanied by alteration in genome organization. Hi-C analysis showed that neither shMcm6 nor *Myc*^ΔORI^ cells underwent major global changes in compartmentalization or topologically associating domain (TADs) integrity (Fig. 4E, F and Fig. S8A, B). Moreover, we detected no discernible topological changes at *Myc* in shMcm6 (Fig. 4G) or *Myc*^ΔORI^ cells (Fig. 4H) or other AID target genes (Fig. S8 C to H). Interestingly, the new ~60 kb late RT domain at *Myc* in *Myc*^ΔORI^ cells spanned approximately equally across either side of the TAD border, which is located ~6 kb upstream of *Myc* (Fig. 4H, lower panel zoom), suggesting that the TAD boundary does not interfere with the establishment of the new late replication domain. In sum, our data show that RT and genome compartmentalization are independent processes and that the decrease in AID-dependent translocations is not due to altered genome architecture. Therefore, we conclude that the loss of AID-dependent translocations is exclusively linked to the loss of early RT at *Myc* in *Myc*^ΔORI^ cells and at AID targets globally in shMcm6 cells

### Re-establishing origin activity and early RT at *Myc* in *Myc*^ΔORI^ cells restores normal *Myc-Igh* translocation frequency

The above results strongly suggest that RT is the only factor regulating the decreased translocation frequency in shMcm6 and *Myc*^ΔORI^ cells. This led us to hypothesize that restoring RT at the *Myc*^ΔORI^ allele would be sufficient to rescue *Myc* translocation frequency. Therefore, we generated a new cell line in which the deleted 800 bp *Myc* origin of replication was inserted into the intergenic space ~14 kb downstream of the *Myc*^ΔORI^ allele (Fig. 5A and Fig. S9 A to C), a region that lies well within the 60 kb late-replicating region created in *Myc*^ΔORI^ cells (Fig. 4D). These new clones, called *Myc*^ΔORI rest^, were viable and proliferative, showed normal *Myc* expression (Fig. S9D) and underwent efficient class switch recombination (Fig. S9 E, F). Repli-seq (Fig. 5B) and Repli-qPCR (Fig. S9G) showed that early RT was fully restored at the *Myc*^ΔORI^ allele in *Myc*^ΔORI rest^ cells suggesting that the *Myc* origin of replication acts autonomously within this replication domain. RT metaplots showed a clear separation of early and late domains genome-wide in *Myc*^ΔORI rest^ cells indicative of a normal RT program, although clone-specific differences were evident (Fig. S9 H, I). Most importantly, *Myc-Igh* translocation frequency was fully reversed to wild-type levels in *Myc*^ΔORI rest^ cells (Fig. 5C). The rescue of *Myc* translocations provides additional confirmation that the parental *Myc*^ΔORI^ lines were not deficient in factors required for translocation but simply lacked early RT at *Myc*. We therefore conclude that the loss of *Myc* translocations in *Myc*^ΔORI^ cells is exclusively due to the loss of origin activity and early RT at *Myc*, thus establishing RT as a critical and distinct step in translocation genesis that directly links DSB formation to DSB ligation.

**Fig. 5.**
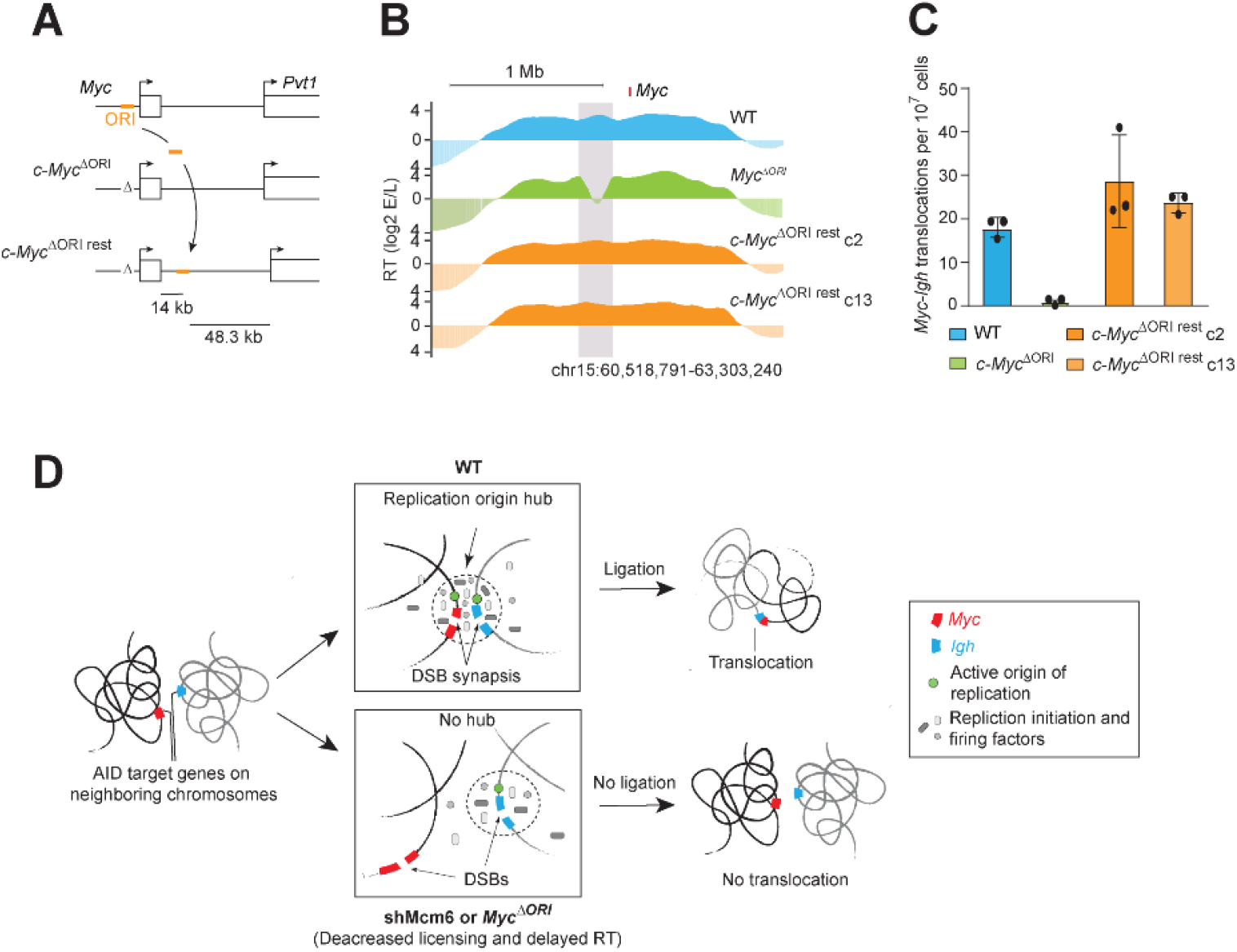
Early RT at *Myc* is essential for translocation genesis and promotes synapsis of DSBs. **(A)** Schematic showing the strategy to restore RT at *Myc* in *Myc*^ΔORI^ cells. The 800 bp origin sequence (orange bar) deleted in *Myc*^ΔORI^ cells (Fig. 2a) was re-inserted 14 kb downstream of *Myc* in *Myc*^ΔORI^ cells and this allele was referred to as *Myc*^ΔORI rest^. **(B)** Snapshot of the RT domains at *Myc* in WT, *Myc*^ΔORI^ and *Myc*^ΔORI rest^ lines showing complete restoration of early RT at the *Myc*^ΔORI rest^ allele compared to *Myc*^ΔORI^. **(C)** Rescue of *Myc-Igh* translocations in *Myc*^ΔORI rest^ lines from three independent experiments. All *P* values in the figure were determined by the unpaired Student’s t test. **(D)** Using *Myc* and *Igh* loci as a paradigm, we propose that a fraction of cells in the population will have both loci in neighboring TADs. In such cells, early origins in *Myc* and *Igh* will occasionally fire synchronously by forming a shared, inter-origin replication hub where pre-RC proteins are concentrated, a concept similar to what has been shown for long-range enhancer-promoter interactions in vivo (*55*). Based on a recent study, we propose that hub formation is seeded via interactions between pre-RC components, ORC, Cdt1 and Cdc6, which can form condensates in vitro that efficiently uptake MCM complexes (*56*). Such a hub would transiently bring DSBs located in the partner loci into close proximity thereby increasing the probability of DSB synapsis. Therefore, if the neighboring origins do not fire synchronously, as in *Myc*^*ΔORI*^ cells or shMcm6 cells, a shared hub is not formed and DSBs in partner loci are less like to interact and create a translocation. Note that since hub formation is only dependent on pre-RC protein-protein interactions and independent of DSBs, the loss of RT synchronicity in *Myc*^*ΔORI*^ or shMcm6 cells cannot be overcome by increasing DSB frequency, as we find in our study. Thus, the RT-mediated step is a distinct regulatory event in translocation genesis independent of DSB frequency.

## Discussion

DNA replication has been proposed to occur in replication hubs or factories wherein multiple origins of replication sharing the same RT congregate and fire simultaneously (*49*). Such hubs are formed via the transient accumulation of protein-protein interactions that can then recruit additional proteins, ultimately leading to a localized region of high protein concentration that facilitate various transactions, such as enhancer-mediated transactivation of gene expression (*55*). In support of this model, ORC, Cdc6, and Cdt1, essential components of the pre-replication complex (pre-RC), were shown to form phase-separated hubs (condensates) in vitro that were able to efficiently recruit MCM complexes for origin activation (*56*). We propose that in cells where AID target genes are located in neighboring chromatin domains, the activation of early replication origins in these genes may occasionally occur synchronously within a shared, inter-origin hub composed of the essential replication proteins (Fig. 5D). Such a process would bring DSBs located near the origins into physical proximity and promote translocation (Fig. 5D). Importantly, hub formation would be driven by interactions between replication factors independently of DSBs (*56*). Therefore, if origin activation of the partner loci is not synchronous (such as in *Myc*^*ΔORI*^ cells or shMcm6 cells), such inter-origin hubs would not form and DSBs would interact less often resulting in fewer translocations (Fig. 5D). The residual translocations seen in *Myc*^*ΔORI*^ cells and shMcm6 cells may occur via random interactions of *trans* DSBs between neighboring TADs and reflect the baseline frequency of translocation. We note that since AID-dependent DSBs are ligated via non-homologous end-joining DNA repair pathways, which are inefficient as cells progress into S phase, DSB ligation would need to occur in early S phase coinciding with the RT of AID target loci.

In summary, we describe here a new and unprecedented step in translocation genesis regulated by the activity of early replication origins and early RT that, we propose, promotes DSB synapsis and thereby physically links DSB formation to DSB ligation. Intriguingly, translocation breakpoints in several cancers have been correlated with early RT and transcriptional activity (*57–60*), but whether early RT was causal for translocation genesis was unknown. We anticipate that our findings will provide the conceptual and experimental framework for a deeper exploration of the role of replication timing in different cancer contexts and towards targeted therapeutic intervention aimed at mitigating oncogenic translocations.

## Acknowledgements

We are grateful to the Vienna Biocenter Core Facilities (VBCF) for next-generation sequencing, recombinant Cre protein and sgRNA synthesis, and the IMP/IMBA core facilities especially the animal house, bio-optics, molecular biology service and proteomics service. We thank Philipp Rescheneder (Oxford Nanopore Technologies), Dominik Handler (IMBA) and Lukas Leindecker (IMP) for valuable input on setting up the MinION platform and supply of essential reagents. We are grateful to Anton Goloborodko (IMBA) and Roman Stocsits (IMP) for advice on Hi-C data analysis. We thank Meinrad Busslinger (IMP), Alex Stark (IMP), Daniel Gerlich (IMBA), Clemens Plaschka (IMP) and Iain Patten for critical reading of the manuscript. This work was funded by Boehringer Ingelheim, The Austrian Industrial Research Promotion Agency (Headquarter Grant FFG-834223), and grant a from the Austrian Science Fund to RP (FWF P 29163-B26).

## Author contributions

MP designed and performed translocation assays, SNS-seq, SNS-qPCR, RNA-seq, LAM-HTGTS, Repli-qPCR, Repli-seq and Hi-C, generated and characterized all the cell lines, and analyzed the data. TN performed bioinformatic analyses of SNS-seq, RNA-seq, LAM-HTGTS, Repli-seq and Hi-C. DM performed bioinformatic analyses of SNS-seq and Hi-C. TN and DM together established the bioinformatic workflows used in this study available at https://github.com/pavrilab/. MN prepared material for nuclear mass spectrometry. US performed PRO-cap. RP conceived and supervised the project, performed ChIP-qPCR, SNS-qPCR and translocation assays, analyzed data and wrote the manuscript.

## MATERIALS AND METHODS

### Cell culture

CH12 cells were maintained in RPMI medium with 10% fetal bovine serum (FBS; Invitrogen), glutamine (Invitrogen), sodium pyruvate (Invitrogen), Hepes (made in-house) and antibiotic/antimycotic (Invitrogen). LentiX and PlatE packaging cells were maintained in DMEM medium with 10% FBS (Invitrogen) and Penicillin/Streptomycin (Invitrogen).

### Mice

Mature, naïve B cells were isolated from spleens of 8-16 week old WT C57BL/6J mice, as per established protocols (*61*). B cells were cultured in complete RPMI medium supplemented with 10% fetal bovine serum and antibiotics, Interleukin 4 (IL4; made in-house by the Molecular Biology Service, IMP) and 25 μg/ml Lipopolysaccharide (Sigma). All animal experiments were carried out with a valid breeding license (No.: GZ: MA58-320337-2019-9) obtained from the Austrian Veterinary Authorities and in compliance with IMP-IMBA animal house regulations.

### CH12 cell activation and class switch recombination (CSR) assay

CH12 cells were activated with 5 ng/mL interleukin-4 (IL-4; home-made), anti-CD40 (home-made), and 1 ng/mL transforming growth factor β (TGF-β; R&D Systems), as described (*43*). For CSR, cells were analyzed 48-72 h later by flow cytometry (BD Fortessa) following staining with anti-IgA coupled to phycoerythrin (PE) (Southern Biotech) (Table S2).

### Transfections and infections

Transfection of LentiX cells with shRNAs against LacZ and Mcm6 (pLKO lentiviral vector TRCN0000124142) followed by infection of CH12 cells with lentiviral supernatants was done exactly as described (*33, 43*). In brief, 24 h after spin-infection (2,350 rpm for 90 min), cells were selected with 1 μg/ml Puromycin (Roche) and activated with IL-4, anti-CD40 and TGF-β 24 h later in the presence of Puromycin. Cells were harvested 48 h after activation. For AIDER expression, the pMX retroviral vector expressing AIDER-IRES-mCherry was transfected into PlatE cells and the viral supernatants (48 h or 72 h) were used to spin-infect CH12 cells for 90 min at 2,350 rpm. AIDER-positive cells were sorted based on mCherry expression. These cells were then infected with shRNAs and activated, as described above. To trigger AIDER nuclear import, 2 μM 4-hydroxy tamoxifen (4-HT; Sigma) was added at the time of activation.

### Cell proliferation assay with Cell Trace Violet dye dilution

The assay was performed using the CellTrace Violet Cell Proliferation Kit (Invitrogen). Cells were stained as per the kit manual and dye content was measured by flow cytometry at 0 h, 24 h and 48 h after staining.

### BrdU incorporation assay for DNA synthesis

Following infection with shRNAs and selection with Puromycin, 100 μM BrdU (Sigma) was added for 48 h followed by staining with anti-BrdU-APC antibody (BD Biosciences) and flow cytometric analysis.

### Genomic DNA extraction

50-200 million cells were washed twice with cold PBS containing 2% FBS and resuspended in 1 ml Proteinase K buffer (100 mM Tris pH 8, 0.2% SDS, 5 mM EDTA, 200 mM NaCl) with 200 μg Proteinase K (Roche). Lysis was performed overnight at 55°C, followed by phenol extraction (Invitrogen UltraPure Phenol:Chloroform:Isoamyl alcohol 25:24:1, v/v), precipitation with ethanol, and resuspension in TE buffer (10 mM Tris pH 8, 1 mM EDTA).

### *Myc-Igh t*ranslocation PCR assay

Nested translocation PCR assays was performed from genomic DNA as previously described (*29*) using either LA Taq DNA polymerase (Takara) or Flex HS polymerase (NEB). The primers are listed in Table S2. PCR products were visualized on an agarose gel, individual bands were excised and cloned into the pCR4 TOPO-TA vector (Invitrogen) or pCR Blunt II-TOPO vector (Invitrogen). Following bacterial transformation, colonies were subjected to plasmid miniprep (Machery Nagel) and isolated plasmids analyzed by Sanger sequencing (Molecular Biology Service, IMP). Where indicated, 1μM Aphidicolin (Sigma) was added at the time of activation.

### Total RNA-seq

For the library generation, 200 ng of high-quality RNA (RIN 9-10) was used per replicate and three biological replicates per condition were processed. Ribosomal RNA removal was done using RiboCop rRNA Depletion Kits (Human/Mouse/Rat) (Lexogen) followed by total RNA library preparation with the CORALL Total RNA-Seq Library Prep Kit (Lexogen) following the manufacturer’s protocols.

### Nuclear proteomics analysis

Nuclei were prepared by resuspending cells in cold sucrose buffer (0.32 M sucrose, 3 mM CaCl2, 2 mM Mg-acetate, 0.1 mM EDTA, 10 mM Tris-HCl (pH 8.0), 0.5% NP-40 and 1 mM DTT). Following centrifugation at 700 *g* and 4°C, the pelleted nuclei were washed in cold sucrose buffer. Nuclei were lysed and subjected to tandem mass tag (TMT) mass spectrometric analysis at the IMP/IMBA Protein Chemistry facility. A complete protocol is available upon request.

### Isolation of short nascent strands (SNSs)

SNSs were isolated following an established protocol kindly provided by Dr. Maria Gomez (CBMSO, Madrid) (*44*) and also described in our previous study (*43*). In brief, 200 million cells per replicate from asynchronous cell cultures were harvested and genomic DNA was extracted. DNA was denatured at 95oC for 10 minutes and then subjected to size fractionation via 5-20% neutral sucrose gradient centrifugation (24,000 *g* for 20 h). Fractions were analyzed by alkaline agarose gel electrophoresis and fractions in the 500-2000 nt range were pooled. Prior to all following enzymatic treatments, ssDNA was heat denatured for 5 min at 95oC. DNA was phosphorylated for 1 h at 37oC with T4 Polynucleotide kinase (NEB or made in-house by the Molecular Biology Service, IMP). To enrich for nascent DNA strands, the phosphorylated ssDNA was digested overnight at 37oC with Lambda exonuclease (NEB or made in-house by the Molecular Biology Service, IMP). Both T4 PNK and Lambda exonuclease steps were repeated twice for a total of three rounds of phosphorylation and digestion. After the final round of digestion, the DNA was treated with RNaseA/T1 mix (Thermo Scientific) to remove 5’ RNA primers and genomic RNA contamination. DNA was purified via phenol-chloroform extraction and ethanol precipitation This material was either used directly for qPCR or further processed for library preparation.

### SNS-qPCR

One-sixth of the SNS preparation (from above) was used for qPCR assays which were performed with the Promega GoTaq master mix (Promega). Each qPCR reaction was carried out in triplicate and in all cases two independent biological replicates per condition were assayed per run. Efficiency of the primers used for qPCR was pre-checked on sonicated DNA. Only primers with the same Cq value or a maximum difference of 1 cycle were used for SNS-qPCR. Primers are listed in Table S2.

### SNS library preparation for SNS-seq

SNSs prepared as described above were converted to double-stranded DNA (dsDNA) via random priming with random hexamer primer phosphate (Roche) and ligation with Taq DNA ligase (NEB). DNA was checked on a fragment analyzer and 50 ng was used for library preparation with the NEBNext® Ultra™ II DNA Library Prep Kit (NEB) following the manufacturer’s protocol. Libraries were barcoded using the NEBNext® Singleplex Oligos (NEB) as per the NEB protocol which allowed 4-8 libraries to be pooled per run. Sequencing was performed on an Illumina HiSeq 2500 machine (50 bp, single-end).

### PRO-cap

PRO-cap in CH12 cells was performed following an established protocol (*62*) with modifications described in detail in our previous study (*61*).

### Generation of *Myc*^ΔORI^ and *Myc*^ΔORI rest^ lines

To delete the *Myc* origin of replication, we generated homology repair plasmids having Ef1a promoter-driven floxed GFP and mCherry expression cassettes flanked by homology arms spanning the region 792 bp and 985 bp on the 5′ and 3′ sides, respectively, of the *Myc* origin. These plasmids were transfected into CH12 cells along with three in vitro synthesized guide RNAs (6 μg each, designed using CRISPOR) and 7.5 μg recombinant Cas9 protein (Vienna Biocenter Core Facilities) using the Neon Transfection System (Thermo Fisher Scientific). A week later, GFP/mCherry double-positive single cells were isolated using a BD FACSAria III sorter (BD Biosciences). Successful knock-in clones were identified by genotyping with PCR and Sanger sequencing of PCR products. Next, 200 μg recombinant Cre recombinase (Molecular Biology Service, IMP) was added to the culture medium to excise the floxed mCherry/GFP cassettes. GFP/mCherry double-negative clones were isolated one week after electroporation using a BD FACSAria III sorter and genotyped via PCR and Sanger sequencing of PCR products. The *Myc*^ΔORI rest^ lines were generated with the same dual-fluorophore approach. Sequences of sgRNAs and primers are listed in Table S2.

### Linear amplification-mediated high-throughput genome-wide translocation capture sequencing (LAM-HTGTS)

The protocol was modified and adapted for the needs of this study from the original protocol (*47*). In brief, 40 million cells per replicate were activated and 12-20 h later electroporated with the pSpCas9(BB)-2A-GFP plasmid (pX458; Addgene 48138) expressing the *Myc* sgRNA using the MaxCyte STx transfection system (MaxCyte). 48 h post-electroporation, GFP-positive cells were collected on a Sony SH800S Cell Sorter followed by genomic DNA extraction. For each sample, eight rounds of linear amplification PCR reactions with 40μg genomic DNA were performed with 5 μg genomic DNA each using HS Flex polymerase (NEB) and biotinylated *Myc* primer (Table S2). PCR reactions were pooled and incubated with 60 μl of Dynabeads™ MyOne™ Streptavidin C1 (Invitrogen) for 1 h at room temperature. DNA was eluted from the beads via phenol extraction or heating in water at 80oC for 5 min and ssDNA adapter [Phos]CCACGCGTGCCCTATAGTCGC[Ami] was ligated using T4 RNA ligase (NEB). Libraries were amplified and barcoded via nested and tagged PCR following the original protocol (*47*) with modified primers (Table S2). Libraries were sequenced using the MiniOn sequencing platform (Oxford Nanopore) following the manufacturer’s protocol. For each sample, a minimum of 1 million sequenced reads were generated.

### Sample preparation for replication timing analysis (Repli-qPCR and Repli-seq)

Repli-seq was performed as previously described (*51, 52*). In brief, two million asynchronously dividing cells were seeded and incubated with 100 μM BrdU (Sigma) for 2 h in a light protected environment to maintain BrdU stability. Cells were fixed and incubated with a mix of RNase A (Invitrogen) and propidium iodide (Sigma) for 30 min (light protected). For each sample, three fractions were sorted: G1 phase, early S phase and late S phase cells, and for each fraction, two independent samples of 50,000 cells (technical replicates) were sorted on a Sony SH800S Cell Sorter. Sorted cells were lysed with Proteinase K buffer overnight. Extracted DNA was sonicated for 9 min in a Diagenode Bioruptor resulting in 100-500 bp DNA fragments as determined on an agarose gel. Sonicated DNA was subjected to end-repair and adapter ligation using the NEBNext® Ultra™ II DNA Library Prep Kit (NEB) following the NEB protocol. Adapter-ligated DNA was incubated with 25 μg/ml of anti-BrdU antibody (BD Pharmingen) for 4h with rotation followed by incubation with 40 μg of anti-mouse IgG antibody (Sigma) for 1 h with rotation (light protected). DNA was precipitated via centrifugation at 16,000 *g* for 5 min at 4°C. The pellet was resuspended in 200 μl of digestion buffer (50 mM Tris-HCl, pH 8.0, 10 mM EDTA and 0.5% SDS) with freshly added 0.25 mg/ml Proteinase K and incubated overnight at 37°C. DNA was purified and used for Repli-qPCR or next generation sequencing (Repli-seq).

### Repli-qPCR

One-sixth of the library DNA (prepared as described above) was used for qPCR. In each experiment, two independent biological replicates per condition were assayed. For each replicate, two G1, two early S and two late S phase fractions were collected and treated as technical replicates. Each qPCR reaction was performed in triplicate. Previously described primers for conserved early and late regions (*51*) were used as controls to assess the quality of the experiment (Table S2).

### Repli-seq

Libraries were barcoded using the NEBNext® Singleplex Oligos (NEB) as per the NEB protocol. Libraries that were successfully validated by Repli-qPCR (as described above) were sequenced on an Illumina HiSeq 2500 machine (50 bp, single-end). Up to 12 barcoded samples were pooled per lane.

### In situ Hi-C

Hi-C was performed as previously described (*54*) with minor modifications. In brief, 5 million cells were crosslinked with 1% formaldehyde (Sigma) for 10 min and quenched with 0.6 M glycine for 5 min. Cells were lysed with Hi-C lysis buffer for 1 h on ice and nuclei were collected by centrifugation. Nuclei were digested with 375 U of Mbol (NEB) overnight at 37oC with rotation. Biotin-14-dATP (Life Technologies) was incorporated for 1 h at 37oC with rotation. Ligation of overhangs was performed with 20,000 U of T4 DNA ligase (NEB) for 4 h at room temperature with rotation. Nuclei were pelleted and reverse-crosslinked overnight. Purified DNA was sonicated for 14 min in a Diagenode Bioruptor to obtain a size range of 200-700 bp. This material was purified using Agencourt AMPure XP beads (Beckman Coulter). Between 8-15 μg of DNA was incubated with 100 μl (~10 mg) of Dynabeads MyOne Streptavidin C1 (Invitrogen) for 15 min with rotation. End repair and adapter ligation using the NEBNext® Ultra™ II DNA Library Prep Kit was performed on-beads following the kit manual. The adapter-ligated DNA was washed, eluted and PCR-amplified with KAPA 2X HiFi HotStart ReadyMix (Kapa Biosystems) and the NEBNext Multiplex Oligos for Illumina® (Dual Index Primers Set 1). Four pooled, barcoded samples were sequenced on an Illumina NovaSeq 6000 machine (50 bp, paired-end).

### Biological and technical replicates

For biological replicates, different frozen vials of CH12 cells were thawed and kept separate throughout the course of the experiment. Replicates for all next-generation sequencing experiments were derived in this manner. Technical replicates, where used (such as in RT-qPCR, Repli-qPCR and SNS-qPCR), were subsets of the biological replicate.

### Statistical analyses

For correlation scatter plots, the Pearson coefficient was calculated to determine the degree of correlation. In all other cases, the two-tailed, unpaired Student’s t test was used for statistical significance.

### Primers, probes, sgRNAs and antibodies

Details of all primers, sgRNAs, probes and antibodies are listed in Table S2.

## Bioinformatics

A detailed methods section for bioinformatics analyses is provided as a supplemental methods file

