## Supplemental figures and methods for "DNA replication timing directly regulates the frequency of oncogenic chromosomal translocations"

#### Supplementary materials

- Supplemental figure legends
- Supplemental figures S1-S9
- Supplemental methods (Bioinformatics)
- Supplemental Tables S1 and S2.

**Supplemental figures S1 to S9**

**Fig. S1: RNA-seq and nuclear mass spectrometry from in shLacZ and shMcm6 cells**

**(A)** Western blot analysis from nuclear extracts of activated shLacZ and shMcm6 CH12 cells. Shown are samples from three experiments with 20 μg extract loaded per well and γ-Tubulin used as a loading control. **(B)** Cell proliferation analysis using CellTrace Violet dye dilution. Cells were stained following shRNA infection and selection and assayed at the indicated time points post-staining. **(C)** DNA synthesis analysis based on incorporation of BrdU. Cells were pulsed with BrdU after infection and selection and assayed 48 h later using an anti-BrdU antibody. **(D)** RNA-seq replicate correlation analysis between triplicate samples of shLacZ and shMcm6 CH12 cells. The color of each square corresponds to the Pearson correlation coefficient of the read counts which is indicated in the color gradient below the plot. **(E)** Gene expression in TPM of *Igh, Myc* and the six MCM complex subunits. **(F)** Protein abundance (normalized peptide counts) from nuclear mass spectrometry for AID, Myc and the MCM complex subunits in shLacZ and shMcm6 cells. **(G)** Scheme of the *Myc-Igh* translocation PCR assay [1] used in the study. **(H) and (I),** Mutation analysis from activated shLacZ and shMcm6 CH12 cells at the translocation hotspot in *Myc* (H) and *Igh* (I) The red bar indicates the sequenced region. *P* values were calculated with the Student's t test.

**Fig. S2: SNS-seq analysis in shLacZ and shMcm6 cells**

**(A)** Workflow of SNS-seq and SNS-qPCR. The assay works on the principle of isolating size-selected genomic DNA fragments followed by two rounds of digestion with λ-exonuclease. The RNA primer at the 5` end of origin-associated SNSs protects them from exonuclease action resulting in enrichment of SNSs over the genomic background. **(B)** Pearson correlation analysis for the SNS-seq replicates. Two independent experiments with two replicates each were performed for shLacZ and shMcm6. **(C)** A representative snapshot of SNS-seq profiles from two independent experiments (exp 1 and exp 2) in shLacZ cells illustrating the calling of initiation sites (red bars) and initiation zones (black bars). For all downstream analyses, only initiation sites within initiation zones were considered (retained initiation sites). A detailed explanation of the workflow is provided in the Methods. **(D) to (E)** Scatter plots of initiation sites are the same as in Fig. 1h but with highlighting of initiation sites overlapping selected histone modifications (D)) or Hi-C-based chromosome subcompartments (E). The numbers of overlapping initiation sites and percentage of overlap are indicated at the bottom right of each plot.

Note how downregulated initiation sites overlap mostly with the activating marks (H3K27ac, H3K4me1, H3K4me3 and H3K36me3) and active A subcompartments whereas the upregulated sites overlap strongly with the silent B2 subcompartment and are enriched for H3K9me3, the mark for constitutive heterochromatin. **(F)** Permutation statistical test with 10,000 draws from translocation hotspot and initiation site-matched null overlaps (grey distribution with the black line indicating the mean) compared to the actual (experimental) overlap between hotspots and initiation sites (green line). The red line represents the statistical significance boundary ($\alpha$ = 0.05). The *P* value is derived from the z-score of the actual observed overlap between hotspots and initiation sites and reflects the difference between the average random overlap following permutations ($OVLP_{perm}$) and the actual observed overlap ($OVLP_{obs}$). **(G)** SNS-qPCR analysis at *Myc*. The top panel shows the SNS-seq track in WT CH12 cells with the location of the amplicons (1-4) used for SNS-qPCR. The bottom panel shows the bar graphs of SNS-qPCR from five experiments in CH12 cells (left) and three experiments in primary, splenic B cells (right). **(H)** SNS-qPCR analysis at *Igh*. The top panel shows the SNS-seq track in WT CH12 cells with the location of the amplicons (1-3) used for SNS-qPCR. The bottom panel shows the bar graphs of SNS-qPCR from three experiments in CH12 cells (left) and three experiments in primary, splenic B cells (right). All *P* values in the above figures were determined by the unpaired Student's t test.

**Fig S3: Generation and characterization of *Myc*$^{\Delta ORI}$ cells**

**(A)** Strategy for deletion of the *Myc* origin of replication (see also Fig. 2a). Recombinant Cas9 and in vitro-synthesized sgRNAs were transfected along with two homology repair template plasmids bearing the same homology arms (HA) but with a central, floxed expression cassette containing either GFP or Cherry. Following successful, homozygous integration, cells were transfected with recombinant Cre protein to excise the floxed cassette yielding the final *Myc*$^{\Delta ORI}$ allele. **(B)** Flow cytometry analysis to generate *Myc*$^{\Delta ORI}$ cells following the scheme in (A). Single cell-sorted clones of GFP/mCherry double-positive cells (gated population in the top row) were expanded (middle row) and genotyped. Correctly integrated clones were transfected with Cre protein, sorted for the absence of both GFP and mCherry (gated double-negative population in the bottom row), and genotyped. **(C)** Genotyping PCR analysis of four, independent *Myc*$^{\Delta ORI}$ clones relative to WT cells. Primer locations are indicated with arrows in the lowest panel in (A) above. **(D)** Proliferation assay using cell trace violet dye dilution analysis. Two *Myc*$^{\Delta ORI}$ clones, c13 and c23 were used and flow cytometry performed 0 h, 24 h and 48 h after dye addition. **(E)** RNA-seq analysis of transcripts with TPM>5 in *Myc*$^{\Delta ORI}$ c13 (left plot) and c23 (right plot) cell lines relative to WT cells. MA plots are based on triplicate experiments. The dotted lines indicate a 2-fold change (log2 FC = 1 or -1). Translocation hotspot genes, DNA replication and repair genes are colored as indicated in the key on the right. **(F)** RT-qPCR analysis for *Myc* expression in *Myc*$^{\Delta ORI}$ c13 and c23 clones normalized to the *Drosophila act5* gene. *Drosophila* S2 cells were added to CH12 cells (1:10 ratio) prior to cell lysis. **(G)** Class switch recombination analysis for IgA expression in *Myc*$^{\Delta ORI}$ c13 and 23 clones 48h after activation. **(H)** Quantification of surface IgA expression data from (G) obtained from three, independent experiments. **(I)** Mutation analysis at the *Myc* hotspot in *Myc*$^{\Delta ORI}$ c13 clones. **(J)** Circos plots of LAM-HTGTS data showing raw, unfiltered *Myc* translocation hotspots (prior to calling AID-dependent hotspots) to the indicated chromosomes in WT (top) and *Myc*$^{\Delta ORI}$c13 (bottom) cells. The location of the *Myc* DSB bait is indicated. **(K)** Same as (J) but for WT (top) and *Myc*$^{\Delta ORI}$ c23 (bottom) translocations. **(L)** Same as (J) but for shLacZ (top) and shMcm6 (bottom) translocations. All *P* values in the figure were determined by the unpaired Student's t test.

**Fig. S4. *Myc* translocations at various AID hotpots in WT and *Myc*$^{\Delta ORI}$ c13 cells**

Examples of LAM-HTGTS results in WT and *Myc*$^{\Delta ORI}$ c13 cells at several recurrent AID-dependent translocation hotspot genes reported in the literature [2-5].

**Fig. S5. *Myc* translocations at various AID hotpots in WT and *Myc*$^{\Delta ORI}$ c23 cells**

Examples of LAM-HTGTS results in WT and *Myc*$^{\Delta ORI}$ c23 cells at the same recurrent AID-dependent translocation hotspot genes shown in Fig. S4.

**Fig. S6. *Myc* translocations at various AID hotpots in shLacZ and shMcm6 cells**

Examples of LAM-HTGTS results in shLacZ and shMcm6 cells at the same recurrent AID-dependent translocation hotspot genes shown in Fig. S4, 5.

**Fig. S7. Analysis of replication timing changes in *Myc*$^{\Delta ORI}$ and shMcm6 cells using Repli-seq**

**(A)** Pearson correlation analysis of Repli-seq replicates of shLacZ and shMcm6 cells. Note that early and late fractions are distinct populations in shLacZ cells but have very high correlation in shMcm6 cells, reflecting the abrogation of the RT program in shMcm6 cells. **(B)** Same as in (A) but comparing replicates of WT and *Myc*$^{\Delta ORI}$ c13 Repli-seq samples. In contrast to (A), early fractions are highly correlated between WT and *Myc*$^{\Delta ORI}$ c13 cells and the same is true for the late fractions, indicating that the RT program is comparable between WT and *Myc*$^{\Delta ORI}$ c13 cells. **(C)** Repli-qPCR assay from four experiments showing the shift of RT at *Myc* from early RT in WT cells to late RT in both *Myc*$^{\Delta ORI}$ clones. The boxed areas highlight the Myc_1 and Myc_2 amplicons located in different parts of the *Myc* region*. Hba-a1* and *Pou5f1* were used as early-replicating control regions and *Zfp42* and *Ascl1* were used as late-replicating control regions.

**Fig. S8. Analysis of genome architecture in shMcm6 and *Myc*$^{\Delta ORI}$ cells**

**(A) to (B)** Hi-C matrices of chromosomes 9 (A) and 12 (B) in shLacZ and shMcm6 cells (left) and in WT and *Myc*$^{\Delta ORI}$ cells (right), displayed alongside Hi-C PC1 (compartmentalization) and RT (log2 E/L) tracks. **(C) to (H)** Examples of Hi-C matrices at six selected AID hotspot gene regions - *Mir142* (C), *Bcl2l11* (D), *Aicda* (E), *Pim1* (F), *Ly6e* (G) and *Cd83* (H) - in shLacZ vs shMcm6 cells (left), and WT vs *Myc*$^{\Delta ORI}$ cells (right). The RT (log2 E/L) and Hi-C compartmentalization (PC1) tracks are shown below each matrix.

**Fig. S9. Characterization of *Myc*$^{\Delta ORI\ rest}$cells**

**(A)** Targeting strategy to insert the 800 bp *Myc* orgin of replication downstream of *Myc* in *Myc*$^{\Delta ORI}$ cells. The approach is identical to that used to make *Myc*$^{\Delta ORI}$ lines, as described in Fig. S3A. **(B)** Representative flow cytometric analysis at each step of *Myc*$^{\Delta ORI\ rest}$ line generation as described for *Myc*$^{\Delta ORI\ rest}$ clones in Fig. S3B. **(C)** Genotyping analysis of *Myc*$^{\Delta ORI\ rest}$ cells after Cre-mediated electroporation. Primers used are shown in the lowest panel of (A) above. Three positive clones were identified (c2, c13, c17). **(D)** RT-qPCR analysis of *Myc* mRNA from three independent experiments in WT, *Myc*$^{\Delta ORI}$ and *Myc*$^{\Delta ORI\ rest}$ cells. **(E) and (F)** Flow cytometry analysis for class switch recombination to IgA (E) and summary from four independent experiments (F) in WT, *Myc*$^{\Delta ORI}$ and *Myc*$^{\Delta ORI\ rest}$ cells. *P* values were determined by the unpaired Student’s t test. **(G) to (H)** RT metaplots from Repli-seq analysis showing the absence of major changes in early and late RT domains in both *Myc*$^{\Delta ORI\ rest}$

lines. Plots on the left show genome-wide RT profiles and those on the right show RT at AID hotspots. **(I)** Repli-qPCR from two independent *Myc*$^{\Delta ORI\ rest}$ lines, c2 and c13. The boxed area highlights the Myc_1 and Myc_2 amplicons located in different parts of the *Myc* region*. Hba-a1* and *Pou5f1* were used as early-replicating control regions and *Zfp42* and *Ascl1* were used as late-replicating control regions.

#### SUPPLEMENTARY TABLES

**Supplementary Table S1: List of hotspots and the number of translocations within them as identified from triplicate LAM-HTGTS analysis.**

The table has three tabs corresponding to three experiments shown in Fig. 3: (1) shLacZ vs shMcm6, (2) WT vs *Myc*$^{\Delta ORI}$ c13 and (3) WT vs *Myc*$^{\Delta ORI}$ c23.

**Supplementary Table S2: List of all primers, probes, sgRNAs and antibodies used in this study**

Fig. S1

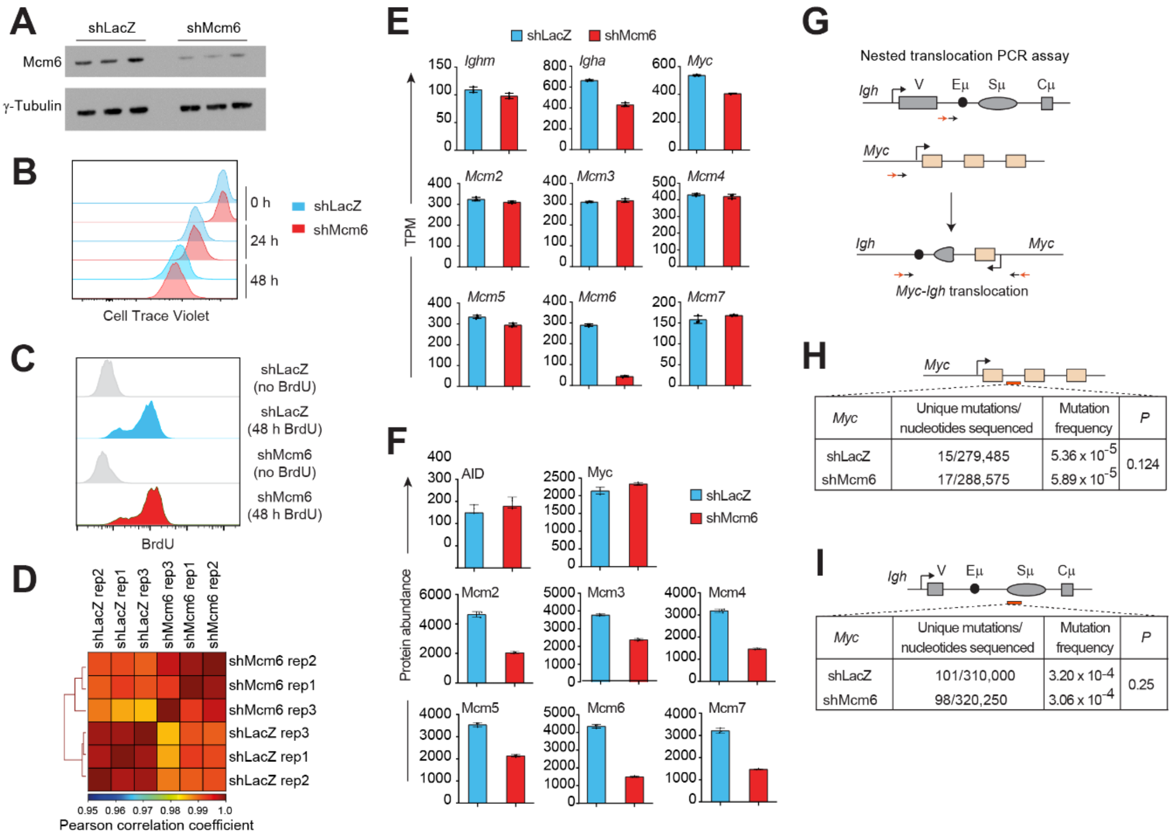


| Myc | Unique mutations/ nucleotides sequenced | Mutation frequency | P |
|---|---|---|---|
| shLacZ | 15/279,485 | $5.36 \times 10^{-5}$ | 0.124 |
| shMcm6 | 17/288,575 | $5.89 \times 10^{-5}$ | |

| Myc | Unique mutations/ nucleotides sequenced | Mutation frequency | P |
|---|---|---|---|
| shLacZ | 101/310,000 | $3.20 \times 10^{-4}$ | 0.25 |
| shMcm6 | 98/320,250 | $3.06 \times 10^{-4}$ | |

Fig. S2

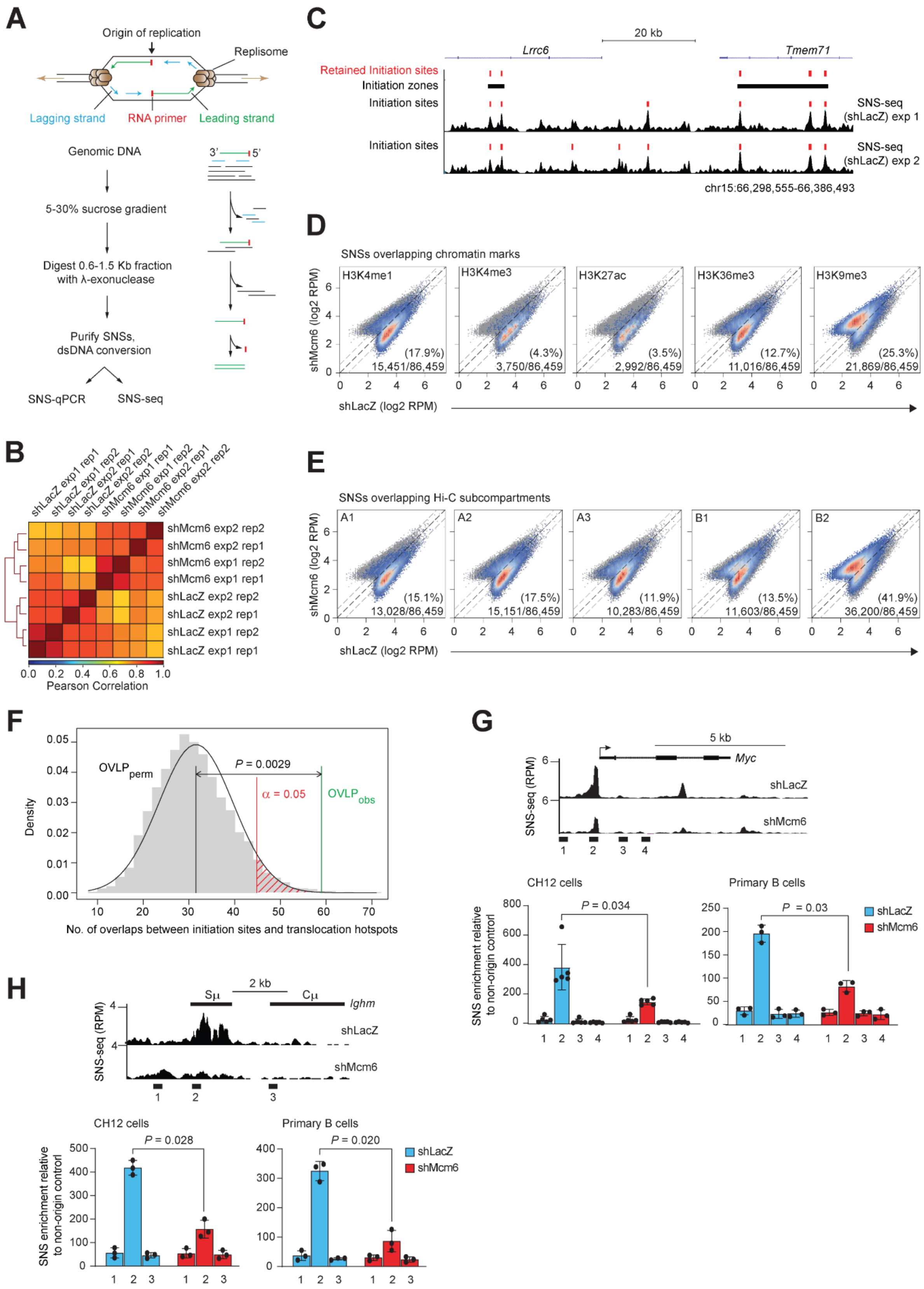
A
Origin of replication
Replisome
Lagging strand
RNA primer
Leading strand
Genomic DNA
5-30% sucrose gradient
Digest 0.6-1.5 Kb fraction with λ-exonuclease
Purify SNSs, dsDNA conversion
SNS-qPCR
SNS-seq
B
shLacZ exp1 rep1
shLacZ exp1 rep2
shLacZ exp2 rep1
shLacZ exp2 rep2
shMcm6 exp1 rep1
shMcm6 exp1 rep2
shMcm6 exp2 rep1
shMcm6 exp2 rep2
Pearson Correlation
C
20 kb
Lrrc6
Tmem71
Retained Initiation sites
Initiation zones
Initiation sites
SNS-seq (shLacZ) exp 1
SNS-seq (shLacZ) exp 2
chr15:66,298,555-66,386,493
D
SNSs overlapping chromatin marks
H3K4me1 (17.9%) 15,451/86,459
H3K4me3 (4.3%) 3,750/86,459
H3K27ac (3.5%) 2,992/86,459
H3K36me3 (12.7%) 11,016/86,459
H3K9me3 (25.3%) 21,869/86,459
shMcm6 (log2 RPM)
shLacZ (log2 RPM)
E
SNSs overlapping Hi-C subcompartments
A1 (15.1%) 13,028/86,459
A2 (17.5%) 15,151/86,459
A3 (11.9%) 10,283/86,459
B1 (13.5%) 11,603/86,459
B2 (41.9%) 36,200/86,459
F
Density
OVLPperm
P = 0.0029
α = 0.05
OVLPobs
No. of overlaps between initiation sites and translocation hotspots
G
5 kb
Myc
SNS-seq (RPM)
shLacZ
shMcm6
CH12 cells
P = 0.034
Primary B cells
P = 0.03
SNS enrichment relative to non-origin control
shLacZ
shMcm6
H
2 kb
Sμ
Cμ
Ighm
SNS-seq (RPM)
shLacZ
shMcm6
CH12 cells
P = 0.028
Primary B cells
P = 0.020
SNS enrichment relative to non-origin control
shLacZ
shMcm6

Fig. S3

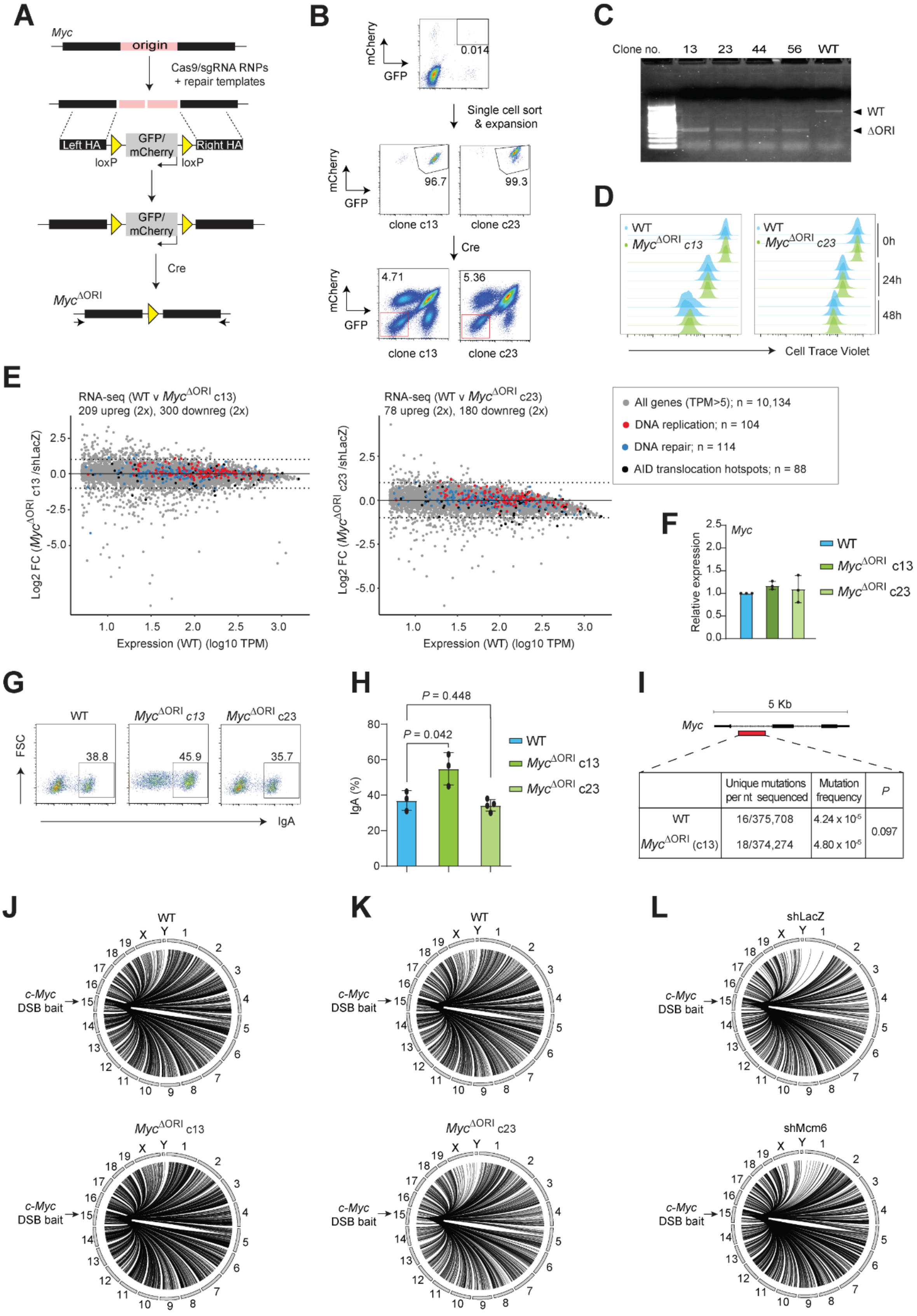
A
Myc
origin
Cas9/sgRNA RNPs
+ repair templates
Left HA
GFP/
mCherry
Right HA
loxP
loxP
GFP/
mCherry
Cre
$Myc^{\Delta ORI}$
B
mCherry
GFP
0.014
Single cell sort
& expansion
96.7
99.3
clone c13
clone c23
Cre
4.71
5.36
clone c13
clone c23
C
Clone no.
13
23
44
56
WT
WT
ΔORI
D
WT
$Myc^{\Delta ORI}$ c13
WT
$Myc^{\Delta ORI}$ c23
0h
24h
48h
Cell Trace Violet
E
RNA-seq (WT v $Myc^{\Delta ORI}$ c13)
209 upreg (2x), 300 downreg (2x)
Log2 FC ($Myc^{\Delta ORI}$ c13 /shLacZ)
Expression (WT) (log10 TPM)
RNA-seq (WT v $Myc^{\Delta ORI}$ c23)
78 upreg (2x), 180 downreg (2x)
Log2 FC ($Myc^{\Delta ORI}$ c23 /shLacZ)
Expression (WT) (log10 TPM)
All genes (TPM>5); n = 10,134
DNA replication; n = 104
DNA repair; n = 114
AID translocation hotspots; n = 88
F
Myc
Relative expression
WT
$Myc^{\Delta ORI}$ c13
$Myc^{\Delta ORI}$ c23
G
WT
$Myc^{\Delta ORI}$ c13
$Myc^{\Delta ORI}$ c23
FSC
38.8
45.9
35.7
IgA
H
P = 0.448
P = 0.042
IgA (%)
WT
$Myc^{\Delta ORI}$ c13
$Myc^{\Delta ORI}$ c23
I
5 Kb
Myc
| | Unique mutations per nt sequenced | Mutation frequency | P |
| WT | 16/375,708 | 4.24 x $10^{-5}$ | 0.097 |
| $Myc^{\Delta ORI}$ (c13) | 18/374,274 | 4.80 x $10^{-5}$ | |
J
WT
$Myc^{\Delta ORI}$ c13
c-Myc
DSB bait
K
WT
$Myc^{\Delta ORI}$ c23
c-Myc
DSB bait
L
shLacZ
shMcm6
c-Myc
DSB bait

### Fig. S4

LAM-HTGTS snapshots of *Myc* translocations to various AID hotspot genes in WT and *Myc*$^{\Delta ORI}$ c13 cells

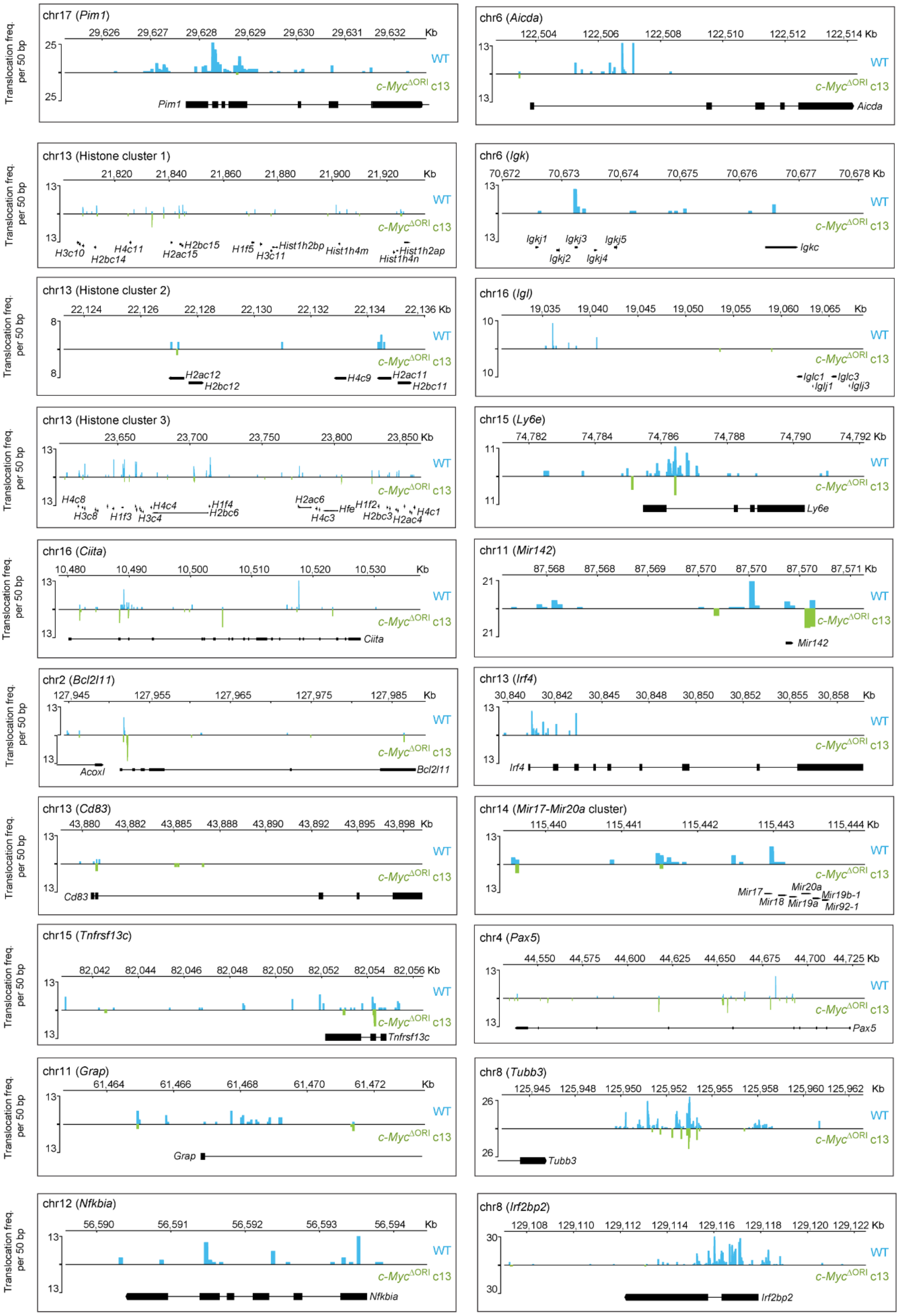

### Fig. S5

LAM-HTGTS snapshots of *Myc* translocations to various AID hotspot genes in WT and *Myc*$^{\Delta ORI}$ c23 cells

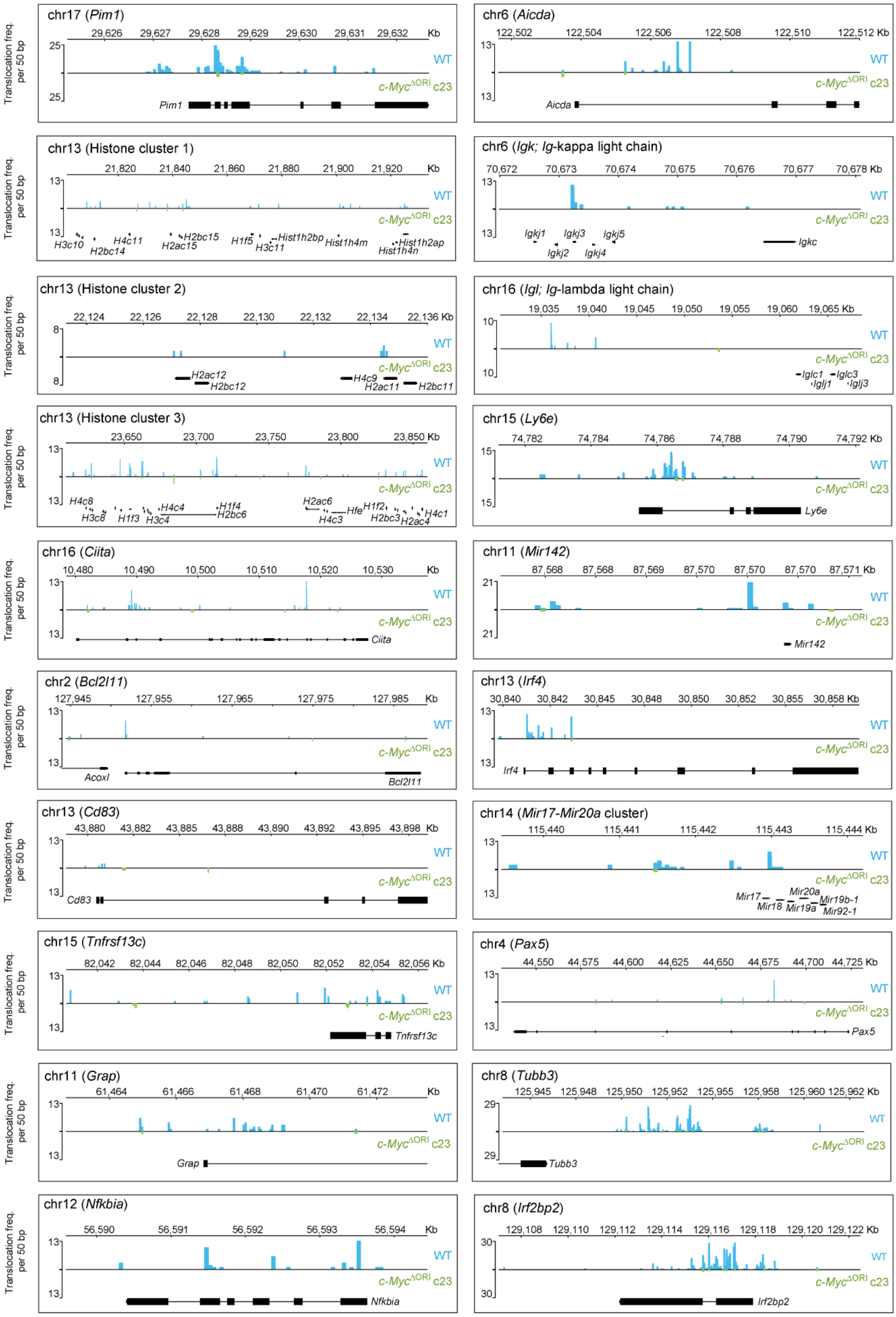

### Fig. S6

LAM-HTGTS snapshots of *Myc* translocations to various AID hotspot genes in shLacZ and shMcm6 cells

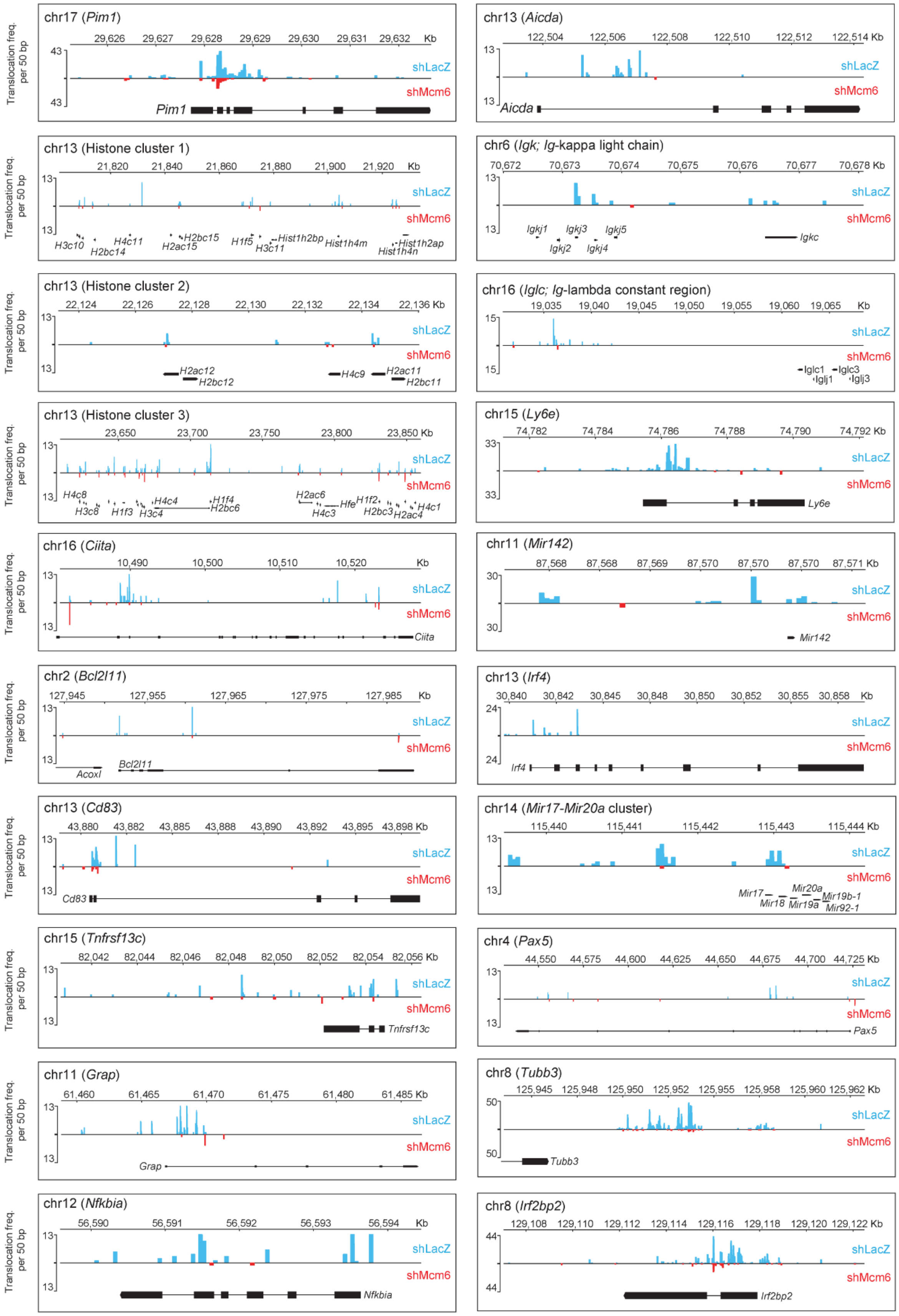

Fig. S7

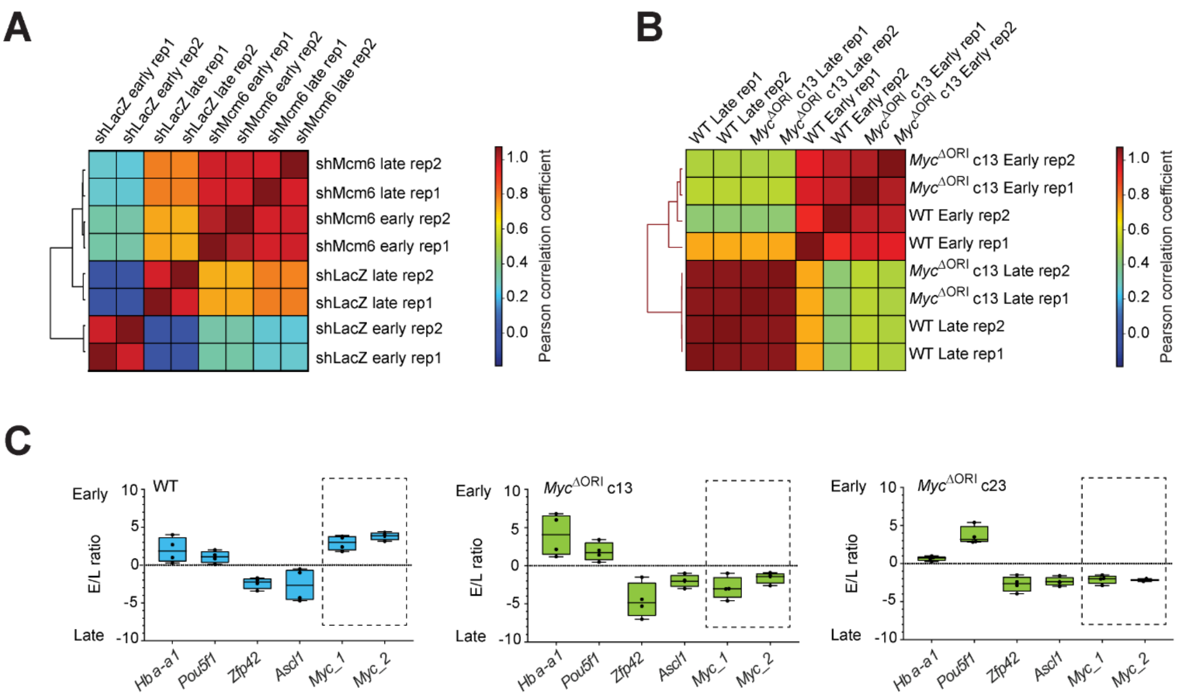
A
shLacZ early rep1
shLacZ early rep2
shLacZ late rep1
shLacZ late rep2
shMcm6 early rep1
shMcm6 early rep2
shMcm6 late rep1
shMcm6 late rep2
shMcm6 late rep2
shMcm6 late rep1
shMcm6 early rep2
shMcm6 early rep1
shLacZ late rep2
shLacZ late rep1
shLacZ early rep2
shLacZ early rep1
Pearson correlation coefficient
1.0
0.8
0.6
0.4
0.2
0.0
B
WT Late rep1
WT Late rep2
MycΔORI c13 Late rep1
MycΔORI c13 Late rep2
WT Early rep1
WT Early rep2
MycΔORI c13 Early rep1
MycΔORI c13 Early rep2
MycΔORI c13 Early rep2
MycΔORI c13 Early rep1
WT Early rep2
WT Early rep1
MycΔORI c13 Late rep2
MycΔORI c13 Late rep1
WT Late rep2
WT Late rep1
Pearson correlation coefficient
C
WT
MycΔORI c13
MycΔORI c23
Early
Late
E/L ratio
10
5
0
-5
-10
Hba-a1
Pou5f1
Zfp42
Ascl1
Myc_1
Myc_2

Fig. S8

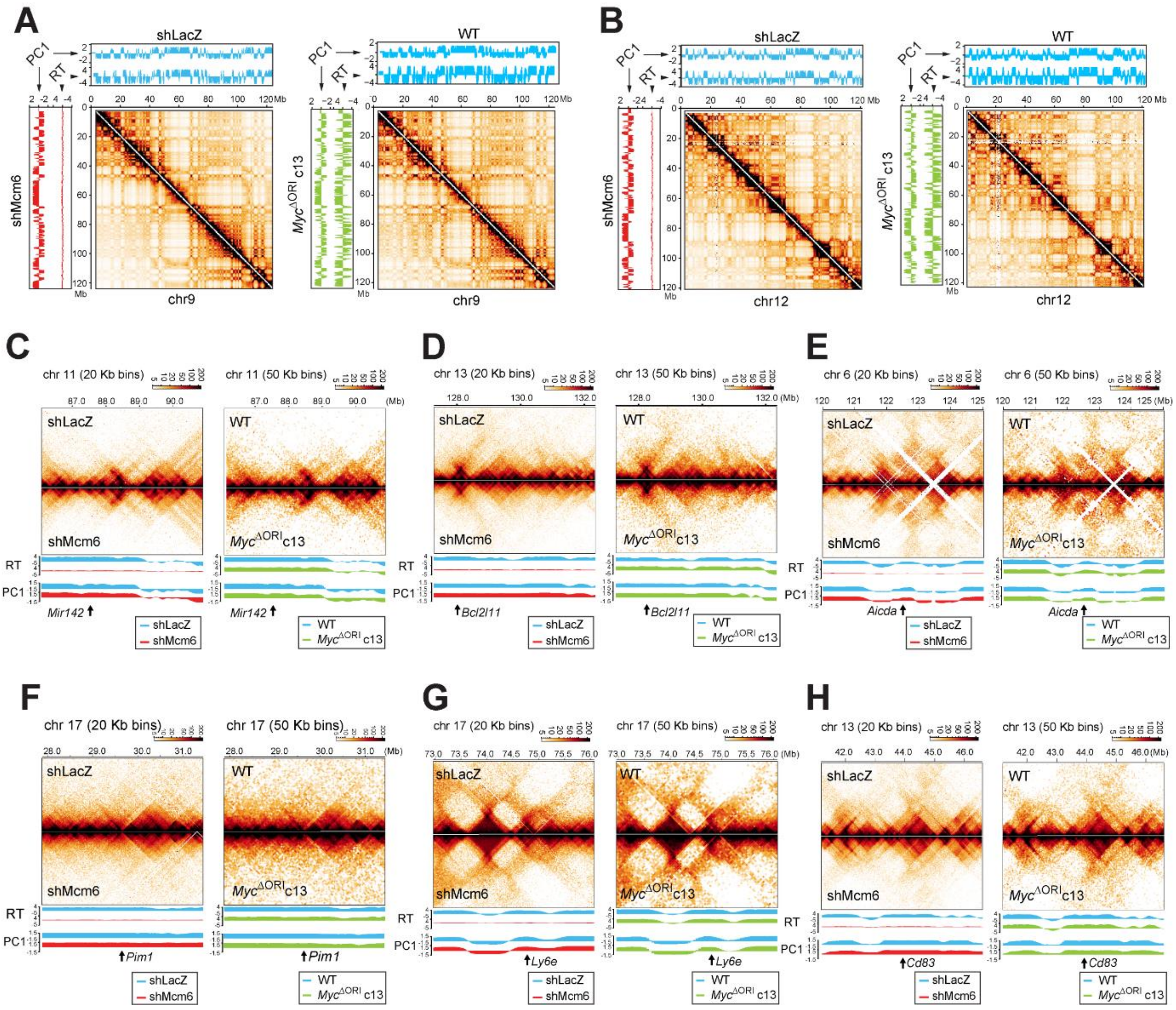
A
B
C
D
E
F
G
H
shLacZ
WT
shMcm6
Myc^ΔORI c13
PC1
RT
chr9
chr12
chr 11 (20 Kb bins)
chr 11 (50 Kb bins)
chr 13 (20 Kb bins)
chr 13 (50 Kb bins)
chr 6 (20 Kb bins)
chr 6 (50 Kb bins)
chr 17 (20 Kb bins)
chr 17 (50 Kb bins)
Mir142
Bcl2l11
Aicda
Pim1
Ly6e
Cd83

Fig. S9

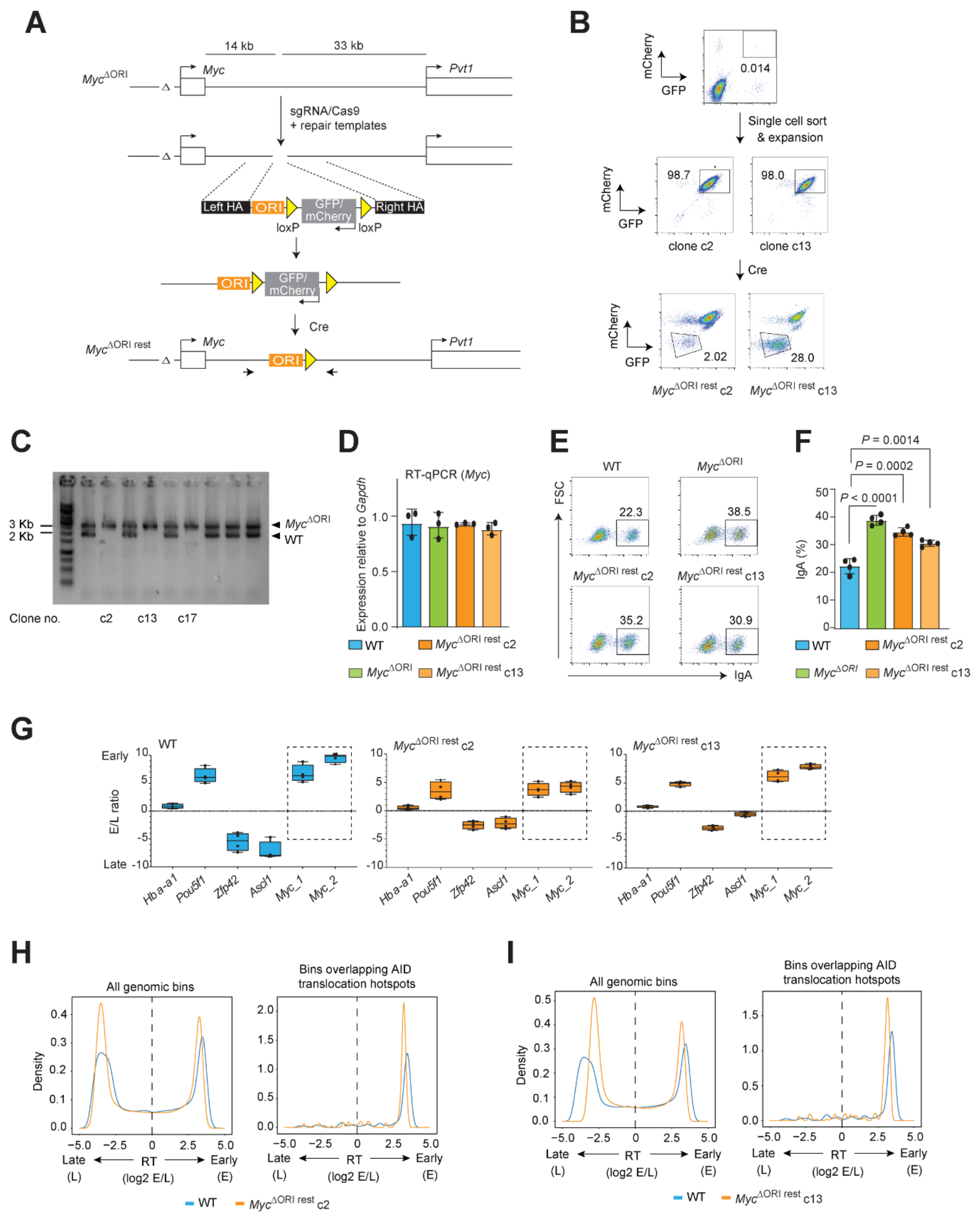

A
14 kb
33 kb
Myc^ΔORI
Myc
Pvt1
sgRNA/Cas9
+ repair templates
Left HA
ORI
GFP/ mCherry
Right HA
loxP
loxP
Cre
Myc^ΔORI rest
B
mCherry
GFP
0.014
Single cell sort & expansion
98.7
98.0
clone c2
clone c13
Cre
2.02
28.0
Myc^ΔORI rest c2
Myc^ΔORI rest c13
C
3 Kb
2 Kb
Myc^ΔORI
WT
Clone no.
c2
c13
c17
D
RT-qPCR (Myc)
Expression relative to Gapdh
WT
Myc^ΔORI
Myc^ΔORI rest c2
Myc^ΔORI rest c13
E
WT
Myc^ΔORI
FSC
22.3
38.5
35.2
30.9
IgA
F
P = 0.0014
P = 0.0002
P < 0.0001
IgA (%)
G
WT
Early
Late
E/L ratio
Hba-a1
Pou5f1
Zfp42
Ascl1
Myc_1
Myc_2
H
All genomic bins
Bins overlapping AID translocation hotspots
Density
Late
(L)
RT
(log2 E/L)
Early
(E)
I

**BIOINFORMATICS**

The established data processing workflows for the main next-generation sequencing datasets produced in this study – SNS-Seq, Repli-Seq and Hi-C - have been developed using the *Nextflow* workflow manager (*1*) and are available centrally at https://github.com/pavrilab. Key features are high-degrees of reproducibility as the compute environment is supplied via Docker and Singularity containers, along with integration of GitHub repositories for self-contained pipelines, portability to all major HPC computing platforms such as SGE, SLURM, AWS, continuous checkpoints for pipeline execution and automatic retrial of failed steps. All workflows produce elaborate QC-reports and out-of-the box resources consumption reports to allow tailoring resources requirements to your datasets which especially for Hi-C datasets can vary by several orders of magnitudes.

**Annotation of initiation sites (ISs) from SNS-seq data**

The analysis workflow was wrapped into the reproducible Nextflow workflow *inisite*-nf available at https://github.com/pavrilab/inisite-nf.

In brief, we performed two independent SNS-seq experiments, each with two technical replicates for WT and KD conditions. Reads were quality and 3' adaptor trimmed with trim_galore (https://github.com/FelixKrueger/TrimGalore) v0.6.4 and cutadapt v2.6 with --match-read-wildcards -O 4 -a AGATCGGAAGAGCACACGTCTGAACTCCAGTCAC and trimmed reads with a length of less than 18bp were discarded. Trimmed reads were then aligned to the mouse genome (mouse genome build UCSC mm9) using bowtie v 1.2.3 (*2*) with -v 2 --best --strata --tryhard -m 1 --chunkmbs 256. The resulting data sets were checked for correspondence between technical replicates by correlation using deeptools multiBamSummary v3.3.0 (*3*). Next, the technical replicates were merged using samtools v1.9 (*4*) to generate a single WT and KD data set per experiment.

Peaks were mapped using MACS2 v2.2.6 (*5*). In particular, we used command-line parameters --nomodel --extsize 275 -q 0.05 and sheared DNA as input. SNS-seq data yielded 95,033 and 119,431 peaks for WT, and 88,578 and 134,649 peaks for KD for experiments 1 and 2, respectively. For each condition, we computed the overlap between the two experiments with bedtools intersect v2.27.1 (*6*) using default parameters. Only peaks found in both experiments were considered for further analysis. Overlapping peaks were merged via bedtools merge v2.27.1 (*6*) using default parameters. This gave 78,627 and 70,621 peaks for WT and KD conditions, respectively.

We used the pandas 0.24.2 Python package (*7*) to compute the median interpeak distance between two neighboring peaks. This value was used to cluster peaks by spatial proximity on the linear genome using ClusterScan v0.2.1 (*8*) setting command-line parameters -n 2 and -d to 16,492 bp and 13,172 kb for WT and KD cells, respectively. Next, we computed the overlap between SNS-seq peaks and clusters with bedtools v2.27.1, only considering peaks lying within a cluster for further analysis. This yielded 56,454 peaks for WT and 47,531 peaks for KD cells. Finally, merging of these peak sets with default (*6*) parameters of bedtools merge v2.27.1(*6*) for each cell type resulted in 86,459 peaks. These peak sets were used for all downstream analyses and are called replication initiation sites (ISs) throughout this study.

**Generation of scatterplots**

Scatterplots were generated using matplotlib v3.1.0 (*9*) by plotting the log2-transformed RPM values for SNS-seq data of the KD against the WT condition computed with deeptools v3.3.0 (*3*). Overlaps are indicated by coloring points using a colormap with the color intensity computed by the number of overlapping points. In particular, a 2D histogram was superposed to the plot area using a 50 x 50 bin grid. Color intensity values were then computed per bin as the density of points in a given bin (i.e. the number of points in a bin divided by the total number of points in the plot). Each point in the given bin is then assigned the color corresponding to the computed density value.

**RNA-seq analysis**

Read alignment

3' adaptors or reads were removed using cutadapt (*10*) v1.4.2 with --match-read-wildcards -O 1 -a AGATCGGAAGAGCACACGTCTGAACTCCAGTCAC) and trimmed reads with a length of less than 18 bp were discarded. The trimmed reads were filtered with a contaminants database consisting of rDNA sequences (gi|374088139, gi|38176281) with bowtie2 v2.1.0 (*11*) --very-sensitive-local. The recovered non matching reads were aligned to the genome/transcriptome using STAR-align v2.4.2a (*12*) with --outSAMstrandField None --outFilterIntronMotifs RemoveNoncanonical --outFilterMismatchNoverLmax 0.1 --outFilterMismatchNmax 10 --outFilterScoreMinOverLread 0.30 --outFilterMatchNminOverLread 0.30 --outFilterMatchNmin 30 --chimSegmentMin 15 --quantMode TranscriptomeSAM --chimJunctionOverhangMin 15 --twopassMode Basic --outSAMtype SAM --outSAMattributes All --outReadsUnmapped Fastx intronMotif --alignIntronMax 200000 --outSJfilterIntronMaxVsReadN 10000 20000 30000 50000 --outSJfilterOverhangMin 20 12 12 12 --outFilterType BySJout --alignMatesGapMax 0 --outFilterMultimapNmax 20. The STAR-index used for alignment was generated from the combination of the mouse genome build NCBI m37 (GCA_000001635.18) and a transcriptome GTF file created from all refGenes and ensGenes gtfs downloaded from the UCSC Table Browser on the 1. March 2014.

Unique mappers from the resulting BAM file were used to create unstranded, stranded and reverse stranded coverage tracks for visualisation using bedtools v2.27 (*6*), bedtools genomecov -split -bg, bedtools genomecov -split -bg -strand '+' ,bedtools genomecov -split -bg -strand '-' and bedGraphToBigWig from the kent-tools bedGraphToBigWig v4 (*13*), bedGraphToBigWig -blockSize=256 -itemsPerSlot=1024. Normalised coverage tracks were created by dividing the coverage at each position with the total number of aligned reads per million.

Differential gene expression analysis: Read counts for genes were quantified using featureCounts v2.0.0 (*14*). As a reference, we downloaded all mm9 refSeq genes from the UCSC table browser (https://hgdownload.soe.ucsc.edu/goldenPath/mm9/bigZips/genes/) on February 28, 2020. In addition, we downloaded the *Igh*, *Igk* and *Igl* gene Ensembl Gene IDs from IMGT (http://www.imgt.org, August 11, 2020) and extracted the corresponding genes from Gencode M25 (www.gencodegenes.org) and lifted them over to mm9 coordinates and added them in addition to the Elambda 3_1 enhancer (10.1101/gad.4.6.978). Differential gene expression analysis was performed with DESeq2 (*15*) v1.22.2 and shrinking log2-foldchanges with ashr v2.2-47 (*16*). To test for unchanged genes, we applied a threshold-based Wald test testing for absolute log-foldChanges < 1.

Gene sets for DNA replication and DNA repair were extracted from the Molecular Signatures Database v7.1 (*17*) using the gene lists listed in the table below.

| **DNA replication** | **DNA repair** |
|---|---|
| GO_CELL_CYCLE_DNA_REPLICATION<br>KEGG_DNA_REPLICATION<br>BIOCARTA_MCM_PATHWAY<br>REACTOME_DNA_REPLICATION_PRE_INITIATION<br>REACTOME_DNA_STRAND_ELONGATION<br>REACTOME_LAGGING_STRAND_SYNTHESIS<br>REACTOME_ACTIVATION_OF_ATR_IN_RESPONSE_TO_REPLICATION_STRESS | KEGG_NON_HOMOLOGOUS_END_JOINING<br>REACTOME_NONHOMOLOGOUS_END_JOINING_NHEJ<br>KEGG_BASE_EXCISION_REPAIR<br>KEGG_MISMATCH_REPAIR<br>REACTOME_DNA_DOUBLE_STRAND_BREAK_REPAIR<br>REACTOME_DNA_DOUBLE_STRAND_BREAK_RESPONSE<br>REACTOME_HOMOLOGOUS_DNA_PAIRING_AND_STRAND_EXCHANGE<br>REACTOME_GLOBAL_GENOME_NUCLEOTIDE_EXCISION_REPAIR_GG_NER<br>REACTOME_HDR_THROUGH_SINGLE_STRAND_ANNEALING_SSA<br>REACTOME_SENSING_OF_DNA_DOUBLE_STRAND_BREAKS<br>REACTOME_PROCESSING_OF_DNA_DOUBLE_STRAND_BREAK_ENDS<br>REACTOME_HDR_THROUGH_HOMOLOGOUS_RECOMBINATION_HRR<br>REACTOME_MISMATCH_REPAIR<br>REACTOME_HOMOLOGY_DIRECTED_REPAIR |

Gene symbols were converted to their mouse homologs using biomaRt v2.38.0 (*18*).

**LAM-HTGTS analysis**

Base-calling and demultiplexing

Raw nanopore fast5 files were basecalled using the *Guppy* basecaller from Oxford Nanopore Technologies (ONT) 3.6.0+98ff765. Resulting reads were demultiplexed by detecting barcodes with a custom fork of *qcat* (https://github.com/nanoporetech/qcat) and splitting reads into chunks at every detected barcode site, assigning the resulting subreads to the majority vote of the detected barcode set. Only reads >= 50 bp were used to remove linker-linker pairs.

Alignment and downsampling

Reads were aligned using bwa mem v0.7.17-r1188 and primary alignments were retained with samtools -F 0x0100. Reads mapping to the Myc bait region (chr15:61794274-61844667) were extracted. Downsampling factors for matched replicates between conditions were calculated and downsampling was performed using picardTools MarkDuplicates (https://broadinstitute.github.io/picard/) v2.18.27.

Translocation and hotspot calling

The *pysam* (https://github.com/pysam-developers/pysam) module was used to extract reads from the *Myc* bait region and translocation breakpoints were extracted from all supplementary alignments in the *SA* bam tag outside the bait region. Translocations also appearing the respective control samples were subtracted for the final translocation sets. Hotspots on the resulting translocation bed files were called using *hot_scan* (*19*) (Version from October 17, 2013) with window width 10 kb, significance level 0.05 and unadjusted p-values.

Confident AID-dependent hotspots

Hotspots from all replicates were overlapped and only hotspots common to all replicates were retained. Hotspots from the WT control sample (which does not express AIDER and is not activated) were subtracted to obtain AID-dependent hotspots. In addition, extragenic hotspots not overlapping with WT RNA Pol II peaks, hotspots < 100 bp in size and low frequency hotspots (< 10 translocations after pooling replicates) were all excluded. This yielded the final set of 88 high-confidence AID-dependent hotspots.

Quantifying changes in translocation frequencies

*Myc* reads from all replicates where taken and pairs of the compared conditions (lacZ vs mcm in the schematic below) were downsampled to equal read numbers. Translocations were called and translocations from a separate control sample were subtracted. The downsampled and control filtered translocation set for each replicate was quantified within the confident hotspots set. Quantifications are pooled between all repliactes and log2 fold-changes between the conditions are calculated by adding a pseudocount of one to avoid division by zero cases.

Association of translocation hotspots with Initiation Sites (IS)

We used the regioneR package to to statistically evaluate the associations between translocation hotspots and Initiation Sites (IS) using permutation tests. 10,000 permutations were performed using the randomizeRegions function on the repeat-masked mouse mm9 genome and overlaps between hotspots and IS were calulcated for each iteration to sample a null distribution. Z-scores and p-values were calculated using the overlapPermTest function.

**Analysis of replication timing (RT) from Repli-seq data**

All RepliSeq analysis was wrapped into the reproducible Nextflow workflow *repliseq-nf* available at https://github.com/pavrilab/repliseq-nf loosely following the steps previously described (*20*). Raw reads from replicates of early (E) and late (L) fractions were trimmed with trim_galore v0.6.5 (https://github.com/FelixKrueger/TrimGalore) and aligned to the mouse mm9 reference sequence using bwa mem v0.7.17 (*21*) and unmapped reads, secondary alignments and multimapping reads were filtered with samtools[6]. Replicates were merged using samtools v1.9 (*4*) and read duplicates were filtered using picardTools MarkDuplicates (https://broadinstitute.github.io/picard/). E/L log2 ratios were calculated in 5kb windows using deeptools bamCompare v3.4.1 (*3*). Loess smoothing was performed with bedtools v2.29.2 (*6*) and a custom R script using the preprocessCore package (https://github.com/bmbolstad/preprocessCore) v1.46.0 with a span size of 300 kb and bigwig tracks were produced using kent_tools v377 (*13*).

**Hi-C analysis**

a. Hi-C data processing

All Hi-C was wrapped into the reproducible Nextflow workflow *hicer*-nf available at https://github.com/pavrilab/hicer-nf. Briefly, raw Hi-C paired-end reads were quality controlled with FastQC v0.11.5 (https://www.bioinformatics.babraham.ac.uk/projects/fastqc/) and raw reads were preprocessed with trim_galore (http://www.bioinformatics.babraham.ac.uk/projects/trim_galore/) with a quality threshold of 20. The reads were then aligned and filtered for experimental artifacts using the HICUP (*22*) pipeline v0.7.3 with bowtie2 v2.3.5.1 (*11*). Resulting SAM files were converted to pairix with linux command line tools and cooler csort (*23*). Pairix files were then used to generate matrices of different resolutions in COOL format using cooler cload (*23*) and cooler zoomify (*23*) and hic format using juicer (*24*). COOL files were balanced with HiCExplorer's (*25*) C++ implementation of the KR algorithm for Python, hic files were balanced with juicer's GW_KR option.

b. Defining A/B compartments

AB compartments were computed with HOMER (*26*) as the first principal component of the Hi-C matrix using a binsize of 20 kb and a smoothing window of 200 kb. Sign correctness of the eigenvector values was established using a genome-wide gene annotation file.

c. Annotation of topologically associating domains and insulation score calculation
Topologically associating domains were called with HOMERs findTADsandLoops.pl (*26*) find mode using a resolution of 5 kb with a 50 kb window and the duplication data for mm9, downloaded from UCSC, as mask for bad regions. The insulation score for subsequent analyses was computed as previously described in Crane et al. 2015n (*27*).

d. Scaling plots and P(s) curves
Scaling plots were computed as the number of ditags separated by a given genomic distance for logarithmically scaled bins from 1 bp to 300 Mb normalized by the total number of valid ditags.

e. Saddle plots
Saddle plots were computed using cooltools v0.3.2. In brief, we subdivided all bins of a 20 kb KR-normalized contact matrix into 50 equal-sized groups based on the bins compartment signal as derived from the eigenvector of the WT data, where group 1 has the lowest signal (i.e. most B) and group 50 has the highest signal (i.e. most A). Subsequently, we compute the mean observed/expected value for each pair of groups and plot it as a 50 x 50 matrix. Similarly, replication timing and H3K9me3 data can be used for bin group assignment.

f. Compartment score computation
The compartment scores in this study were computed as previously described in Nuebler et al. 2018 (*28*).

**PRO-cap analysis**
The Illumina reads were adapter trimmed on their 3′ ends using cutadapt (*10*) v1.15; --match-read-wildcards -f fastq -a AGATCGGAAGAGCACACGTCTGAACTCCAGTCAC). Afterwards, four nucleotides from their 5′ end were removed with fastx_trimmer (http://hannonlab.cshl.edu/fastx_toolkit; version 0.0.13; -Q33 -t 5). The trimmed reads >18 nucleotides were aligned with bowtie v1.0.0 (*2*) (-S -p 10 --trim5 0 --trim3 0 -v 2 --best --strata --tryhard -m 1 --phred33-quals --chunkmbs 256) to the mouse mm9 genome. The aligned reads were sorted by position with samtools v1.9 (*4*). Strand-specific and undirected occupancy profiles were generated with deepTools bamCoverage v2.2.2 (*3*).

**Summary of used Software**
All custom Python scripts and tools, except for scripts using ClusterScan, were developed, tested and executed using Python v3.7.3. ClusterScan and related scripts were executed using Python v2.7.13. Used R-packages were obtained from Bioconductor v3.8 (*29*) via the BiocManager v1.30.10 (https://github.com/Bioconductor/BiocManager) and executed in R v3.5.1 (*30*) in augmentation of the rpy2 programming language interface v3.0.5 (https://rpy2.bitbucket.io/) to use them in conjunction with Python. The used software and packages are summarized in **Error! Reference source not found.**, **Error! Reference source not found.** and **Error! Reference source not found.**. All code is available at https://bitbucket.org/mnmldani/orichoice.

*Table 1: Summary of used standalone software*

| Software | Version | Usage |
|---|---|---|
| bedtools | 2.27.1 | general operations on sets of genomic regions |
| biomaRt | v2.38.0 | Gene symbol conversion |
| bowtie | 1.0.0/1.2.3 | Alignment of reads |
| bowtie2 | 2.2.9 | Alignment of reads |
| bwa | 0.7.17 | Alignment of Repli-seq data |
| ClusterScan | 0.2.1 | Clustering of initiation sites |
| cutadapt | 2.6 | Adapter trimming |
| deeptools | 3.3.0/3.4.1 | Read counting in regions and heatmap generation |
| FastQC | 0.11.5 | Quality control of sequencing reads |

| | | |
|---|---|---|
| featureCounts | 2.0.0 | Computing RNA-seq data counts |
| Guppy | 3.6.0+98ff765 | Nanopore read base calling |
| HICUP | 0.7.3 | Processing of Hi-C data |
| HOMER | 4.10 | Hi-C Eigenvector and TAD annotation |
| hot_scan | Oct 2013 | Annotation of translocation hotspots |
| UCSCkentUtils | 356 | Conversion between file formats |
| MACS2 | 2.2.6 | Peak calling for narrow peak type data |
| Picard Tools | 2.18.27 | BAM Deduplication |
| Python2 | 2.7.13 | Execution of ClusterScan and associated scripts |
| Python3 | 3.7.3 | General script execution |
| R | 3.5.1 | Execution of R code |
| samtools | 1.9 | General operations on aligned reads |
| STAR | 2.4.2a | Alignment of RNA-seq data |
| Trim_galore | 0.6.4 | Quality control of sequencing data |

*Table 2: Summary of used R packages*

| ***Package*** | ***Version*** | ***Usage*** |
|---|---|---|
| ashr | 2.2-47 | Differential expression analysis |
| BiocManager | 1.30.10 | Accessing Bioconductor 3.8 packages |
| DESeq2 | 1.22.2 | Differential expression analysis |

*Table 3: Summary of used Python packages*

| **Package** | **Version** | **Usage** |
|---|---|---|
| argparse | 1.1 | Passing command-line arguments to scripts |
| cooler | 0.8.3 | Hi-C matrix generation and storage |
| cooltools | 0.3.2 | Generic Hi-C downstream analysis |
| krbalancing | 0.0.5 | HiCExplorer implementation of KR for matrix correction |
| logging | 0.5.1.2 | Printing information to stdout |
| matplotlib | 3.1.0 | Generation of general plots |
| numpy | 1.16.4 | General computations on data in matrices |
| pandas | 1.1.0 | General computations on tabulated data |
| pybedtools | 0.8.0 | Using bedtools within Python |
| pyBigWig | 0.3.16 | Generation of bigWig files |
| pysam | 0.15.2 | Using samtools in Python |
| re | 2.2.1 | Regular expressions for efficient text parsing |
| scipy | 1.3.0 | General statistical computation in Python |
| tables | 3.5.2 | Reading of HDF5 files |

#### Methods-only references


1 Di Tommaso, P. *et al.* Nextflow enables reproducible computational workflows. *Nature Biotechnology* **35**, 316-319, doi:10.1038/nbt.3820 (2017).

2 Langmead, B., Trapnell, C., Pop, M. & Salzberg, S. L. Ultrafast and memory-efficient alignment of short DNA sequences to the human genome. *Genome Biology* **10**, R25, doi:10.1186/gb-2009-10-3-r25 (2009).

3 Ramírez, F. *et al.* deepTools2: a next generation web server for deep-sequencing data analysis. *Nucleic Acids Research* **44**, W160-W165, doi:10.1093/nar/gkw257 (2016).

4 Li, H. *et al.* The Sequence Alignment/Map format and SAMtools. *Bioinformatics* **25**, 2078-2079, doi:10.1093/bioinformatics/btp352 (2009).

5 Zhang, Y. *et al.* Model-based Analysis of ChIP-Seq (MACS). *Genome Biology* **9**, R137, doi:10.1186/gb-2008-9-9-r137 (2008).

6 Quinlan, A. R. & Hall, I. M. BEDTools: a flexible suite of utilities for comparing genomic features. *Bioinformatics* **26**, 841-842, doi:10.1093/bioinformatics/btq033 (2010).

7 McKinney, W. Data Structures for Statistical Computing in Python. *Proceedings of the 9th Python in Science Conference*, 51-56 (2010).

8 Volpe, M., Miralto, M., Gustincich, S. & Sanges, R. ClusterScan: simple and generalistic identification of genomic clusters. *Bioinformatics* **34**, 3921-3923, doi:10.1093/bioinformatics/bty486 (2018).
9 Hunter, J. D. Matplotlib: A 2D Graphics Environment. *Computing in Science & Engineering* **9**, 90-95, doi:10.1109/MCSE.2007.55 (2007).
10 Martin, M. Cutadapt removes adapter sequences from high-throughput sequencing reads. *2011* **17**, 3, doi:10.14806/ej.17.1.200 (2011).
11 Langmead, B. & Salzberg, S. L. Fast gapped-read alignment with Bowtie 2. *Nature Methods* **9**, 357-359, doi:10.1038/nmeth.1923 (2012).
12 Dobin, A. *et al.* STAR: ultrafast universal RNA-seq aligner. *Bioinformatics* **29**, 15-21, doi:10.1093/bioinformatics/bts635 (2013).
13 Kent, W. J., Zweig, A. S., Barber, G., Hinrichs, A. S. & Karolchik, D. BigWig and BigBed: enabling browsing of large distributed datasets. *Bioinformatics* **26**, 2204-2207, doi:10.1093/bioinformatics/btq351 (2010).
14 Liao, Y., Smyth, G. K. & Shi, W. featureCounts: an efficient general purpose program for assigning sequence reads to genomic features. *Bioinformatics* **30**, 923-930, doi:10.1093/bioinformatics/btt656 (2013).
15 Love, M. I., Huber, W. & Anders, S. Moderated estimation of fold change and dispersion for RNA-seq data with DESeq2. *Genome biology* **15**, 550-550, doi:10.1186/s13059-014-0550-8 (2014).
16 Stephens, M. False discovery rates: a new deal. *Biostatistics* **18**, 275-294, doi:10.1093/biostatistics/kxw041 (2016).
17 Subramanian, A. *et al.* Gene set enrichment analysis: A knowledge-based approach for interpreting genome-wide expression profiles. *Proceedings of the National Academy of Sciences* **102**, 15545, doi:10.1073/pnas.0506580102 (2005).
18 Durinck, S., Spellman, P. T., Birney, E. & Huber, W. Mapping identifiers for the integration of genomic datasets with the R/Bioconductor package biomaRt. *Nature Protocols* **4**, 1184-1191, doi:10.1038/nprot.2009.97 (2009).
19 Silva, I. T., Rosales, R. A., Holanda, A. J., Nussenzweig, M. C. & Jankovic, M. Identification of chromosomal translocation hotspots via scan statistics. *Bioinformatics* **30**, 2551-2558, doi:10.1093/bioinformatics/btu351 (2014).
20 Marchal, C. *et al.* Genome-wide analysis of replication timing by next-generation sequencing with E/L Repli-seq. *Nature Protocols* **13**, 819, doi:10.1038/nprot.2017.148 (2018).
21 Li, H. & Durbin, R. Fast and accurate short read alignment with Burrows-Wheeler transform. *Bioinformatics* **25**, 1754-1760, doi:10.1093/bioinformatics/btp324 (2009).
22 Wingett, S. *et al.* HiCUP: pipeline for mapping and processing Hi-C data. *F1000Research* **4**, doi:10.12688/f1000research.7334.1 (2015).
23 Abdennur, N. & Mirny, L. A. Cooler: scalable storage for Hi-C data and other genomically labeled arrays. *Bioinformatics* **36**, 311-316, doi:10.1093/bioinformatics/btz540 (2019).
24 Durand, N. C. *et al.* Juicer Provides a One-Click System for Analyzing Loop-Resolution Hi-C Experiments. *Cell Systems* **3**, 95-98, doi:10.1016/j.cels.2016.07.002 (2016).
25 Wolff, J. *et al.* Galaxy HiCExplorer: a web server for reproducible Hi-C data analysis, quality control and visualization. *Nucleic Acids Research* **46**, W11-W16, doi:10.1093/nar/gky504 (2018).
26 Heinz, S. *et al.* Simple combinations of lineage-determining transcription factors prime cis-regulatory elements required for macrophage and B cell identities. *Mol Cell* **38**, 576-589, doi:10.1016/j.molcel.2010.05.004 (2010).
27 Crane, E. *et al.* Condensin-driven remodelling of X chromosome topology during dosage compensation. *Nature* **523**, 240-244, doi:10.1038/nature14450 (2015).
28 Nuebler, J., Fudenberg, G., Imakaev, M., Abdennur, N. & Mirny, L. A. Chromatin organization by an interplay of loop extrusion and compartmental segregation.

*Proceedings of the National Academy of Sciences* **115**, E6697, doi:10.1073/pnas.1717730115 (2018).
29 Huber, W. *et al.* Orchestrating high-throughput genomic analysis with Bioconductor. *Nat Methods* **12**, 115-121, doi:10.1038/nmeth.3252 (2015).
30 RCoreTeam. R: A language and environment for statistical computing. *R Foundation for Statistical Computing* (2018).
