## Supplemental Table iS2 for "DNA replication timing directly regulates the frequency of oncogenic chromosomal translocations"

### Table S2: List of primers, sgRNA, adapters and antibodies

| | | | |
|---|---|---|---|
| **Nested translocation PCR** | | | Source |
| primer set 1 | | | |
| PCR1: Igh | GCAATGACTGAA GACTCAGTCCCTCTTAAG | | Dorsett et al. J. Exp Med 2017 |
| PCR2: Igh | TACCATTTGCGGTGCCTGGTTTCGGAGAGG | | Dorsett et al. J. Exp Med 2017 |
| PCR1: Myc | ACTTAGCCCTGCAGACGCCCAGGAATCGCC | | Dorsett et al. J. Exp Med 2017 |
| PCR2: myc | TTGGCTTCAGAGGCTGAGGGAGGCGACTG | | Dorsett et al. J. Exp Med 2017 |
| primer set 2 | | | |
| PCR1: Igh | GTGCCCCACTCCACTCTTTGTCCCTATGC | | Dorsett et al. J. Exp Med 2017 |
| PCR2: Igh | ATCATCCAGGGACTCCACCAACACCATCAC | | Dorsett et al. J. Exp Med 2017 |
| PCR1: Myc | GAAATAAAAGGGGAGGGGGTGTCAAATAATAAGAG | | Dorsett et al. J. Exp Med 2017 |
| PCR2: myc | CCTCCCTTCTACACTCTAAACCGCGACGCCAC | | Dorsett et al. J. Exp Med 2017 |
| der 15 | | | |
| PCR1: Igh | GTTGAGACATGGGTCTGGGTCAGGGAC | | Dorsett et al. J. Exp Med 2017 |
| PCR2: Igh | CTCTGCCTGCTGGTCTGTGGTGACATTAG | | Dorsett et al. J. Exp Med 2017 |
| PCR1: Myc | ATCAGCGGCCGCAACCCTCGCCGCCGC | | Dorsett et al. J. Exp Med 2017 |
| PCR2: myc | GAAGGCTGGATTTCCTTTGGGCGTTGG | | Dorsett et al. J. Exp Med 2017 |
| der 12 IgH-Smu | | | |
| der 12 Myc PCR2 | GACACCTCCCTTCTACACTCTAAACCG | | Robbiani et al. Cell 2008 |
| der 12 Ig-mu PCR2 | CACCCTGCTATTTCCTTGTTGCTAC | | Robbiani et al. Cell 2008 |
| der 12 Myc PCR1 | GGGGAGGGGGTGTCAAATAATAAGA | | Robbiani et al. Cell 2008 |
| der15 Smu_Igh | | | |
| der 12 Ig-mu PCR1 | TGAGGACCAGAGAGGGATAAAAGAGAA | | Robbiani et al. Cell 2008 |
| der 15 Myc PCR2 | GTGGAGGTGTATGGGGTGTAGAC | | Robbiani et al. Cell 2008 |
| der 15 Ig-mu PCR2 | CCTCAGTCACCGTCTCCTCAGGTA | | Robbiani et al. Cell 2008 |
| der 15 Myc PCR1 | GTGAAAACCGACTGTGGCCCTGGAA | | Robbiani et al. Cell 2008 |
| der 15 Ig-mu PCR1 | ACTATGCTATGGACTACTGGGGTCAAG | | Robbiani et al. Cell 2008 |
| der Salpha-Myc | | | |
| der15 salpa PCR2 | GTGGGATTTCTCGCAGACTCTG | | Ramiro et al. Cell 2004 |
| der15 salpha PCR1 | GAGCTGGTGGGAGTGTCAGTG | | Ramiro et al. Cell 2004 |
| der 15 Myc PCR2 | GTGGAGGTGTATGGGGTGTAGAC | | Ramiro et al. Cell 2004 |
| der 15 Myc PCR1 | GTGAAAACCGACTGTGGCCCTGGAA | | Ramiro et al. Cell 2004 |
| **Sanger sequencing primers for translocation PCR** | | | |
| Smu sequencing primer | CCAATACCCAAAGCATTTACAG | | this study |
| S alpha reverse for seq-correct | CCAGAGGAATACAGGACATGATC | | this study |
| S alpha sequencing | GGATTGCCTTCTTAGCCTGGGTC | | this study |
| S alpha for seq R | GATCATGTCCTGTATTCCTCTGG | | this study |
| S alpha for seq F | CTGATGTCAACCCTGTGACCTC | | this study |
| Myc2_sanger | GACCGGCAGAGACTCCTCCCG | | this study |
| Myc1_sanger | GTATGGGGTGTAGACCGG | | this study |
| Sanger_Ig alpha2 | GGAACATAAAACATGCATGAG | | this study |
| Sanger_Ig alpha1 | CATGAGATAGTAGTCTTCACAC | | this study |
| Sanger_myc2 | CGAACCGGAGGTGCTGGAGT | | this study |
| Sanger_myc1 | CCACACCACCCGCCGGGTGG | | this study |
| fw_myc_mu sanger5 | GCGCTATTACTGTTTACACCCC | | this study |
| rev_myc_mu sanger4 | GCCGCTACATTCAAGACGCA | | this study |
| fw_myc_mu sanger3 | TCCCGAGTTCCCAAAGCAGA | | this study |
| rev_myc_mu sanger2 | CCGGCCGCTACATTCAAGA | | this study |
| fw_myc_mu sanger1 | GCCCTTTATATTCCGGGGGT | | this study |
| 6myc_fw translocseq | GGTTACTATGGGCTGACGCT | | this study |
| 5myc_fw translocseq | GGATGCATTTTGAAGCGGGG | | this study |
| 4myc_sangerseq transloc | TTCTTTGGCCACCACGAAGC | | this study |
| 3myc_sangerseq transloc | CCGGTTTGACCCCTCAAAGG | | this study |
| Salpha translocationseq4 | GGCTCCATTCACACAAAGCAA | | this study |
| Salpha translocationseq3 | CCATGCAGGCACCATTTACAG | | this study |
| Smu translocationseq3 | ACAGTGCTTAGATCCGAGGTG | | this study |
| Smu translocationseq2 | TGAGGTACTGATGCTGTCTCT | | this study |
| Smu translocationseq | TGAGTACTTGAAAACCCTCTCACA | | this study |
| Myc_translocseq2 | GAGCGGGGGCTCCTAGATAA | | this study |
| Myc_translocseq1 | ATCACTCCACACACTGAGCG | | this study |
| **RT-qPCR** | | | |
| Myc | CGCGATCAGCTCTCCTGAAA | GACACCTCCCTTCTACACTCTAAACCG | this study |
| Gapdh | TGAAGCAGGCATCTGAGGG | CGAAGGTGGAAGAGTGGGAG | Fitz et. al. Nat. Genetics 2020 |
| ***Myc* mutation analysis** | | | |
| Myc_mutation pcr | TGGTCTTTCCCTGTGTTCTTTCTG | GACACCTCCCTTCTACACTCTAAACCG | Ramiro et. al. Cell 2004 |
| **SNS qPCR primers** | | | |
| Non-origin region | CCAAACTTCAGCAGTTTTCC | GAAATTGTCTTGCAAGAGGA | this study |
| 1Mycori | CGCGAGCAAGAGAAAATGGTC | GGCTCCGGGGTGTAAACAGT | this study |
| 2Mycori | CGAACAACCGTACAGAAAGGG | ACAGTAATAGCGCAGCATGA | this study |
| 3Mycori | GAGAGGTGGGGAAGGGAGAA | CCCCGGAATATAAAGGGCGG | this study |
| 1mycsns | CAAGGCAAGCGAACAGCAAG | TTCAAACACTGGACAAGGGCT | this study |
| 3mycsns | CAATTCGTAAGCAGTCGGGC | GCATGCTATGTGAGCCCTCT | this study |
| 5mycsns | TGTGTACTTGGTGACGGTCT | GCTAGAGGATCAGGTCACGC | this study |
| 7mycsns | TGAATGTAGCGGCCGGTTAG | CCCGACTCAGATCTACCCCA | this study |
| 9mycsns | GAAGCGAGAGGTTGGATGGA | CACGGGAATCCAGGGATGAAG | this study |
| 11mycsns | GGTGCTTGGGAAATGTGCTTT | GGGTTAGGGCACAGGTGAGA | this study |
| 13mycsns | ATTGTTAAAATGGGGCTGGGG | CACTTGCCTTTTGCCCATTCC | this study |
| 15mycsns | GGTGAAGTGGGAGGAGCTTT | AGGGATCCTCTCGGGGAAAC | this study |
| 17mycsns | TGCCATGGCTAGCTTGGTTT | CGGGAAGTGACGAGAGCAAA | this study |
| Sa Forward | TGAATTGAGCAATGTTGAGTTGGAGTCAAGATG | ACCACTTCTTCAAACCACAGCTACAAGT | Rajagopal et al J Exp Med 2009 |
| Sb Forward | CCCGAAGCATTTACAGTGACTTTGTTCATGA | GATTTGTGAAGCCGTTTTGACCAGAATGTC | Rajagopal et al J Exp Med 2009 |
| Sc Forwar | GGGAATGTATGGTTGTGGCTTCTGC | GTCCAGAGTCTTTGTGTGGAATTGTTCCT | Rajagopal et al J Exp Med 2009 |
| Sd Forward | GTTGCCTGTTAACCAATAATCATAGAGCTCATGG | GTATAACTGAAGTAGAGACAGCATCAGTACCTCAAC | Rajagopal et al J Exp Med 2009 |
| Se Forward, | ACACTACTACATTCTTGATCTACAACTCAATGTGGT | CGGATCTAAGCACTGTCCTTTGATACCATTC | Rajagopal et al J Exp Med 2009 |
| Sf Forward, | CCAGACAGAGAAAGCCAGACTCATAAAGC | GAAGCACTCAGAGAAGCCCACC | Rajagopal et al J Exp Med 2009 |
| Sg Forward, | CTGCCTACACTGGACTGTTCTGAGC | CAGCTCACCCCATCTCACCCCATC | Rajagopal et al J Exp Med 2009 |
| Sh Forward, | CTGTGGTGAGCGGAGCTGGATTGAAC | CTGATCCCAGCACAATCTGGCTCAC | Rajagopal et al J Exp Med 2009 |
| Si Forward, | GTGTAGGGTGATCTGGACTCAACTGG | CTTTTCCCAGCTCATCCCGAACC | Rajagopal et al J Exp Med 2009 |
| Sj Forward | AGCCTGAGCTGAGTAGGTCTAAACTGAG | GAACGAGGTCCTAGCCTAACTGGTC | Rajagopal et al J Exp Med 2009 |
| **LAM-HTGTS** | | | |
| c-Myc-RED-lm | CCTCTGAAGCCAAGGCCGATG | | this sudy |
| c-Myc-Bio-lm | [Btn]GCCTCGGCTCTTAGCAGACTG | | this sudy |
| Adapter oligo | [Phos]CCACGCGTGCCCTATAGTCGC[Ami] | | Hu et. al. Nat. Protocols 2016 |
| Nested PCR | | | |
| barcode1 | CACAAAGACACCGACAACTTTCTTCCTCTGAAGCCAAGGCCGATG | AAGAAAGTTGTCGGTGTCTTTGTGGACTATAGGGCACGCGTGG | this study |
| barcode2 | ACAGACGACTACAAACGGAATCGACCTCTGAAGCCAAGGCCGATG | TCGATTCCGTTTGTAGTCGTCTGTGACTATAGGGCACGCGTGG | this study |
| barcode3 | CCTGGTAACTGGGACACAAGACTCCCTCTGAAGCCAAGGCCGATG | GAGTCTTGTGTCCCAGTTACCAGGGACTATAGGGCACGCGTGG | this study |
| Tagged PCR | | | |
| b1-1 | AAATCTCCTAGATCGCCACAAAGACACCGACAACTTTCTTCCTCTGAAGCCAAGGCCGATG | AAATCTCCTAGATCGCAAGAAAGTTGTCGGTGTCTTTGTGGACTATAGGGCACGCGTGG | this study |
| b1-2 | TTAGTGTACTCTCTATCACAAAGACACCGACAACTTTCTTCCTCTGAAGCCAAGGCCGATG | TTAGTGTACTCTCTATAAGAAAGTTGTCGGTGTCTTTGTGGACTATAGGGCACGCGTGG | this study |
| b1-3 | AGTTCAGATATCCTCTCACAAAGACACCGACAACTTTCTTCCTCTGAAGCCAAGGCCGATG | AGTTCAGATATCCTCTAAGAAAGTTGTCGGTGTCTTTGTGGACTATAGGGCACGCGTGG | this study |
| b1-4 | CCAATTGGAGAGTAGACACAAAGACACCGACAACTTTCTTCCTCTGAAGCCAAGGCCGATG | CCAATTGGAGAGTAGAAAGAAAGTTGTCGGTGTCTTTGTGGACTATAGGGCACGCGTGG | this study |
| b1-5 | TAGTTTGTGTAAGGAGCACAAAGACACCGACAACTTTCTTCCTCTGAAGCCAAGGCCGATG | TAGTTTGTGTAAGGAGAAGAAAGTTGTCGGTGTCTTTGTGGACTATAGGGCACGCGTGG | this study |
| b1-6 | CCAGAACTACTGCATACACAAAGACACCGACAACTTTCTTCCTCTGAAGCCAAGGCCGATG | CCAGAACTACTGCATAAAGAAAGTTGTCGGTGTCTTTGTGGACTATAGGGCACGCGTGG | this study |
| b1-7 | CAGATTTTAAGGAGTACACAAAGACACCGACAACTTTCTTCCTCTGAAGCCAAGGCCGATG | CAGATTTTAAGGAGTAAAGAAAGTTGTCGGTGTCTTTGTGGACTATAGGGCACGCGTGG | this study |
| b1-8 | AACAGCAGTAAGGCGACACAAAGACACCGACAACTTTCTTCCTCTGAAGCCAAGGCCGATG | AACAGCAGTAAGGCGAAAGAAAGTTGTCGGTGTCTTTGTGGACTATAGGGCACGCGTGG | this study |
| b1-9 | AGGACGCACGTACTAGCACAAAGACACCGACAACTTTCTTCCTCTGAAGCCAAGGCCGATG | AGGACGCACGTACTAGAAGAAAGTTGTCGGTGTCTTTGTGGACTATAGGGCACGCGTGG | this study |
| b1-10 | TGCTCTACAGGCAGAACACAAAGACACCGACAACTTTCTTCCTCTGAAGCCAAGGCCGATG | TGCTCTACAGGCAGAAAAGAAAGTTGTCGGTGTCTTTGTGGACTATAGGGCACGCGTGG | this study |
| b2-1 | AAATCTCCTAGATCGCACAGACGACTACAAACGGAATCGACCTCTGAAGCCAAGGCCGATG | AAATCTCCTAGATCGCTCGATTCCGTTTGTAGTCGTCTGTGACTATAGGGCACGCGTGG | this study |
| b2-2 | TTAGTGTACTCTCTATACAGACGACTACAAACGGAATCGACCTCTGAAGCCAAGGCCGATG | TTAGTGTACTCTCTATTCGATTCCGTTTGTAGTCGTCTGTGACTATAGGGCACGCGTGG | this study |
| b2-3 | AGTTCAGATATCCTCTACAGACGACTACAAACGGAATCGACCTCTGAAGCCAAGGCCGATG | AGTTCAGATATCCTCTTCGATTCCGTTTGTAGTCGTCTGTGACTATAGGGCACGCGTGG | this study |
| b2-4 | CCAATTGGAGAGTAGAACAGACGACTACAAACGGAATCGACCTCTGAAGCCAAGGCCGATG | CCAATTGGAGAGTAGATCGATTCCGTTTGTAGTCGTCTGTGACTATAGGGCACGCGTGG | this study |
| b2-5 | TAGTTTGTGTAAGGAGACAGACGACTACAAACGGAATCGACCTCTGAAGCCAAGGCCGATG | TAGTTTGTGTAAGGAGTCGATTCCGTTTGTAGTCGTCTGTGACTATAGGGCACGCGTGG | this study |
| b2-6 | CCAGAACTACTGCATAACAGACGACTACAAACGGAATCGACCTCTGAAGCCAAGGCCGATG | CCAGAACTACTGCATATCGATTCCGTTTGTAGTCGTCTGTGACTATAGGGCACGCGTGG | this study |
| b2-7 | CAGATTTTAAGGAGTAACAGACGACTACAAACGGAATCGACCTCTGAAGCCAAGGCCGATG | CAGATTTTAAGGAGTATCGATTCCGTTTGTAGTCGTCTGTGACTATAGGGCACGCGTGG | this study |
| b2-8 | AACAGCAGTAAGGCGAACAGACGACTACAAACGGAATCGACCTCTGAAGCCAAGGCCGATG | AACAGCAGTAAGGCGATCGATTCCGTTTGTAGTCGTCTGTGACTATAGGGCACGCGTGG | this study |
| b2-9 | AGGACGCACGTACTAGACAGACGACTACAAACGGAATCGACCTCTGAAGCCAAGGCCGATG | AGGACGCACGTACTAGTCGATTCCGTTTGTAGTCGTCTGTGACTATAGGGCACGCGTGG | this study |
| b2-10 | TGCTCTACAGGCAGAAACAGACGACTACAAACGGAATCGACCTCTGAAGCCAAGGCCGATG | TGCTCTACAGGCAGAATCGATTCCGTTTGTAGTCGTCTGTGACTATAGGGCACGCGTGG | this study |
| b3-1 | AAATCTCCTAGATCGCCCTGGTAACTGGGACACAAGACTCCCTCTGAAGCCAAGGCCGATG | AAATCTCCTAGATCGCGAGTCTTGTGTCCCAGTTACCAGGGACTATAGGGCACGCGTGG | this study |
| b3-2 | TTAGTGTACTCTCTATCCTGGTAACTGGGACACAAGACTCCCTCTGAAGCCAAGGCCGATG | TTAGTGTACTCTCTATGAGTCTTGTGTCCCAGTTACCAGGGACTATAGGGCACGCGTGG | this study |
| b3-3 | AGTTCAGATATCCTCTCCTGGTAACTGGGACACAAGACTCCCTCTGAAGCCAAGGCCGATG | AGTTCAGATATCCTCTGAGTCTTGTGTCCCAGTTACCAGGGACTATAGGGCACGCGTGG | this study |
| b3-4 | CCAATTGGAGAGTAGACCTGGTAACTGGGACACAAGACTCCCTCTGAAGCCAAGGCCGATG | CCAATTGGAGAGTAGAGAGTCTTGTGTCCCAGTTACCAGGGACTATAGGGCACGCGTGG | this study |
| b3-5 | TAGTTTGTGTAAGGAGCCTGGTAACTGGGACACAAGACTCCCTCTGAAGCCAAGGCCGATG | TAGTTTGTGTAAGGAGGAGTCTTGTGTCCCAGTTACCAGGGACTATAGGGCACGCGTGG | this study |
| b3-6 | CCAGAACTACTGCATACCTGGTAACTGGGACACAAGACTCCCTCTGAAGCCAAGGCCGATG | CCAGAACTACTGCATAGAGTCTTGTGTCCCAGTTACCAGGGACTATAGGGCACGCGTGG | this study |
| b3-7 | CAGATTTTAAGGAGTACCTGGTAACTGGGACACAAGACTCCCTCTGAAGCCAAGGCCGATG | CAGATTTTAAGGAGTAGAGTCTTGTGTCCCAGTTACCAGGGACTATAGGGCACGCGTGG | this study |
| b3-8 | AACAGCAGTAAGGCGACCTGGTAACTGGGACACAAGACTCCCTCTGAAGCCAAGGCCGATG | AACAGCAGTAAGGCGAGAGTCTTGTGTCCCAGTTACCAGGGACTATAGGGCACGCGTGG | this study |
| b3-9 | AGGACGCACGTACTAGCCTGGTAACTGGGACACAAGACTCCCTCTGAAGCCAAGGCCGATG | AGGACGCACGTACTAGGAGTCTTGTGTCCCAGTTACCAGGGACTATAGGGCACGCGTGG | this study |
| b3-10 | TGCTCTACAGGCAGAACCTGGTAACTGGGACACAAGACTCCCTCTGAAGCCAAGGCCGATG | TGCTCTACAGGCAGAAGAGTCTTGTGTCCCAGTTACCAGGGACTATAGGGCACGCGTGG | this study |
| **NEB Indexing primers** | | | |
| Index primers | sequence | expected index primer read | Source |
| Index primer set1 | | | |
| Index primer 1 | 5´-CAAGCAGAAGACGGCATACGAGATCGTGATGTGACTGGAGTTCAGACGTGTGCTCTTCCGATC-s-T-3´ | ATCACG | NEB |
| Index primer 2 | 5´-CAAGCAGAAGACGGCATACGAGATACATCGGTGACTGGAGTTCAGACGTGTGCTCTTCCGATC-s-T-3´ | CGATGT | NEB |
| Index primer 3 | 5´-CAAGCAGAAGACGGCATACGAGATGCCTAAGTGACTGGAGTTCAGACGTGTGCTCTTCCGATC-s-T-3´ | TTAGGC | NEB |
| Index primer 4 | 5´-CAAGCAGAAGACGGCATACGAGATTGGTCAGTGACTGGAGTTCAGACGTGTGCTCTTCCGATC-s-T-3´ | TGACCA | NEB |
| Index primer 5 | 5´-CAAGCAGAAGACGGCATACGAGATCACTGTGTGACTGGAGTTCAGACGTGTGCTCTTCCGATC-s-T-3´ | ACAGTG | NEB |
| Index primer 6 | 5´-CAAGCAGAAGACGGCATACGAGATATTGGCGTGACTGGAGTTCAGACGTGTGCTCTTCCGATC-s-T-3´ | GCCAAT | NEB |
| Index primer 7 | 5´-CAAGCAGAAGACGGCATACGAGATGATCTGGTGACTGGAGTTCAGACGTGTGCTCTTCCGATC-s-T-3 | CAGATC | NEB |
| Index primer 8 | 5´-CAAGCAGAAGACGGCATACGAGATTCAAGTGTGACTGGAGTTCAGACGTGTGCTCTTCCGATC-s-T-3´ | ACTTGA | NEB |

| | | | |
|---|---|---|---|
| Index primer 9 | 5´-CAAGCAGAAGACGGCATACGAGATCTGATCGTGACTGGAGTTCAGACGTGTGCTCTTCCGATC-s-T-3´ | GATCAG | NEB |
| Index primer 10 | 5´-CAAGCAGAAGACGGCATACGAGATAAGCTAGTGACTGGAGTTCAGACGTGTGCTCTTCCGATC-s-T-3´ | TAGCTT | NEB |
| Index primer 11 | 5´-CAAGCAGAAGACGGCATACGAGATGTAGCCGTGACTGGAGTTCAGACGTGTGCTCTTCCGATC-s-T-3´ | GGCTAC | NEB |
| Index primer 12 | 5´-CAAGCAGAAGACGGCATACGAGATTACAAGGTGACTGGAGTTCAGACGTGTGCTCTTCCGATC-s-T-3´ | CTTGTA | NEB |
| NEBNext Adaptor for Illumina | 5´-/5Phos/GATCGGAAGAGCACACGTCTGAACTC AGTCdU CACTCTTTCCCTACACGACGCTCTTCCGATC-s-T-3´ | N/A | NEB |
| NEBNext Universal PCR Primer for Illum | 5´-AAT GAT ACG GCG ACC ACC GAG ATC TAC ACT CTT TCC CTA CAC GAC GCT CTT CCG ATC-s-T-3´ | N/A | NEB |
| Dual index primer set1 | | | |
| NEBNext i501 Primer | 5´-AATGATACGGCGACCACCGAGATCTACACTATAGCCTACACTCTTTCCCTACACGACGCTCTTCCGATC*T-3´ | TATAGCCT | NEB |
| NEBNext i502 Primer | 5´-AATGATACGGCGACCACCGAGATCTACACATAGAGGCACACTCTTTCCCTACACGACGCTCTTCCGATC*T-3´ | ATAGAGGC | NEB |
| NEBNext i503 Primer | 5´-AATGATACGGCGACCACCGAGATCTACACCCTATCCTACACTCTTTCCCTACACGACGCTCTTCCGATC*T-3´ | CCTATCCT | NEB |
| NEBNext i504 Primer | 5´-AATGATACGGCGACCACCGAGATCTACACGGCTCTGAACACTCTTTCCCTACACGACGCTCTTCCGATC*T-3´ | GGCTCTGA | NEB |
| NEBNext i701 Primer | 5´-CAAGCAGAAGACGGCATACGAGATCGAGTAATGTGACTGGAGTTCAGACGTGTGCTCTTCCGATC*T-3´ | ATTACTCG | NEB |
| NEBNext i702 Primer | 5´-CAAGCAGAAGACGGCATACGAGATTCTCCGGAGTGACTGGAGTTCAGACGTGTGCTCTTCCGATC*T-3´ | TCCGGAGA | NEB |
| NEBNext i703 Primer | 5´-CAAGCAGAAGACGGCATACGAGATAATGAGCGGTGACTGGAGTTCAGACGTGTGCTCTTCCGATC*T-3´ | CGCTCATT | NEB |
| NEBNext i704 Primer | 5´-CAAGCAGAAGACGGCATACGAGATGGAATCTCGTGACTGGAGTTCAGACGTGTGCTCTTCCGATC*T-3´ | GAGATTCC | NEB |
| **Repli qPCR** | | | |
| mt DNA | GACATCTGGTTCTTACTTCA | GTTTTTGGGGTTTGGCATTA | Ryba et al. Nat. Protocols 2011 |
| Hbaa1 | AAGGGGAGCAGAGGCATCA | AGGGCTTGGGAGGGACTG | Ryba et al. Nat. Protocols 2011 |
| Hbbb1 | CAGTAAGCCACAGATCCTATTG | CCCATAGTGACTATTGACTGTG | Ryba et al. Nat. Protocols 2011 |
| Pou5 | CCCTCCCTAAGTGCCAGTTTCT | GTAATCGCCCTCAGCAGTGTCT | Ryba et al. Nat. Protocols 2011 |
| Mmp15 | AACAGAAGGCCTGCCTTGAC | TGCATAGCACGACAGCATTG | Ryba et al. Nat. Protocols 2011 |
| Zfp42 | TGAGATTAGCCCCGAGACTGAG | CGTCCCCTTTGTCATGTACTCC | Ryba et al. Nat. Protocols 2011 |
| Mash1 | GAAGATGAGCAAGGTGGAGACG | AGTAGGACGAGACCGGAGAACC | Ryba et al. Nat. Protocols 2011 |
| Akt3 | GAAGTGTGGGTTGAACCTCTGG | GCACCCTCTCCACTGTTCTGAT | Ryba et al. Nat. Protocols 2011 |
| Myc1 | CGCGAGCAAGAGAAAATGGTC | GGCTCCGGGGTGTAAACAGT | this study |
| Myc2 | CGAACAACCGTACAGAAAGGG | ACAGTAATAGCGCAGCATGA | this study |
| Myc3 | GAGAGGTGGGGAAGGGAGAA | CCCCGGAATATAAAGGGCGG | this study |
| **c-Myc ΔORI lines genotyping** | | | |
| Myc_after Cre | TCCCTTCTTTTTCCCGCCAA | CTTGTGAAAACCGACTGTGGC | this study |
| GFP | ACAATGAGCACCTTATACACGC | CCCGACAACCACTACCTGAG | this study |
| PCR_myc_sffvprom | GCGGTGACCATCTGTTCTTG | CAGGCTGTCAGAAATGCACC | this study |
| mCherry | TCCTAGAGGCTGTAGTCATTTTGC | CCTTCAGCTTGGCGGTCTG | this study |
| Myc origin | ATACACAATGCCTTCCCCGC | GCTGGAATTACTACAGCGAGTC | this study |
| **sgRNA sequences** | | | |
| sgRNA for LAM-HTGTS | GACGAGCGTCACTGATAGTA | | Panchakshari et al. PNAS 2018 |
| sgRNA for MycΔORI 1 | AGCCGGAGTACTGGGCTGCG | | this study |
| sgRNA for MycΔORI 2 | CGGTTCGGACTTCCACCCGG | | this study |
| sgRNA for MycΔORI rest 1 | CGCAGTGGTCATGTAACT | | this study |
| sgRNA for MycΔORI rest 2 | GTCATGTAACTCGGGCGACA | | this study |
| sgRNA for MycΔORI rest 3 | ATGTAACTCGGGCGACATGG | | this study |
| **Antibodies** | | | |
| | Manufacturer | catalog number | dillution |
| FACS | | | |
| IgA-PE | Southern Biotech | SB-1040-09 | 1/500 for 1x10^6 cells |
| anti-BrdU-APC | BD Pharmingen | 51-2319L/2300525 | 10 µM |
| Western blot | | | |
| Mcm6 | Santa Cruz (H-8) | sc-393618 | 1/500 |
| beta-Tubulin | Sigma (TUB2.1) | T-5201 | 1/2000 |
| ChIP-qPCR | | | |
| Mcm5 | Bethyl Laboratories | A300-195A | 10 µg for 20 million cells |
